# Immiscible proteins compete for RNA binding to order condensate layers

**DOI:** 10.1101/2025.03.18.644007

**Authors:** Wilton T. Snead, Mary K. Skillicorn, Krishna Shrinivas, Amy S. Gladfelter

**Affiliations:** Department of Cell Biology, Duke University School of Medicine, Durham, NC, USA; Department of Chemical and Biological Engineering, Northwestern University, Evanston, IL, USA

**Keywords:** biomolecular condensates, RNA, RNA-binding proteins, nuclear paraspeckles

## Abstract

Biomolecular condensates mediate diverse and essential cellular functions by compartmentalizing biochemical pathways. Many condensates have internal subdomains with distinct compositional identities. A major challenge lies in dissecting the multicomponent logic that relates biomolecular features to emergent condensate organization. Nuclear paraspeckles are paradigmatic examples of multi-domain condensates, comprising core and shell layers with distinct compositions that are scaffolded by the lncRNA NEAT1, which spans both layers. A prevailing model of paraspeckle assembly proposes that core proteins bind directly and specifically to core-associated NEAT1 domains. Combining informatics and biochemistry, we unexpectedly find that the essential core proteins FUS and NONO bind and condense preferentially with shell-associated NEAT1 domains. The shell protein TDP-43 exhibits similar NEAT1 domain preferences on its own but forms surfactant-like shell layers around core protein-driven condensates when both are present. Together, experiments and physics-based simulations suggest that competitive RNA binding and immiscibility between core and shell proteins orders paraspeckle layers. More generally, we propose that sub-condensate organization can spontaneously arise from a balance of collaborative and competitive protein binding to the same domains of a lncRNA.

**Significance:** The cellular milieu is spatially organized into compartments called biomolecular condensates that exhibit rich internal organization which shapes their functions. Often comprising multiple proteins and RNAs, a major question concerns how molecular-scale features relate to emergent condensate forms. Here we study nuclear paraspeckles, archetypal multi-domain condensates comprising distinct core and shell layers assembled around a layer-spanning RNA scaffold. We find that different proteins associated with each layer all bind preferentially to the same shell-associated domains of the RNA scaffold. Core and shell proteins are inherently immiscible, establishing a competition that redirects core proteins to a suboptimal, core-associated domain of the scaffold. Our work reveals how a combination of competitive RNA binding and protein immiscibility can spatially organize multicomponent condensate subdomains.

## Introduction

Biomolecular condensation is an important type of emergent physical organization in living cells. Distinct nuclear condensates regulate essential functions that span transcription^1^, splicing^2,3^, chromatin organization^4,5^, and ribosome biogenesis^6^ – in large part by recruiting complex yet specific molecular communities. Unlike a well-mixed compartment of molecules, many condensates exhibit intricate and spatially varying architectures, compositions, and physical properties. In prominent examples, spanning nucleoli^6^, nuclear speckles^7^, stress granules^8^, and germ granules in *C. elegans*^9^ and *D. melanogaster*^10^, disruption of sub-condensate composition and architecture leads to dysregulated downstream function. A key challenge in condensate biology lies in decoding the mechanisms by which multicomponent biomolecular networks confer condensates with distinct identities and spatially varying architectures, and thereby drive their emergent functions.

Nuclear paraspeckles are paradigmatic examples of multicomponent condensates composed of core and shell layers with distinct compositions^11,12^. Specifically, the RNA-binding proteins (RBPs) FUS and NONO are localized in the core and TDP-43 in the shell^13,14^. The long noncoding RNA (lncRNA) NEAT1 is the central layer-spanning scaffold that recruits RBPs and specifies layer identity^15,16^. Paraspeckles regulate post-transcriptional gene expression by controlling nuclear-cytoplasm export of specific mRNAs^17–19^. In turn, paraspeckles impact a variety of cellular processes in both health and disease^12^, including immune cell activation^20^, lineage specification in early development^21^, and cancer cell fitness and motility^22,23^. Importantly, the core-shell arrangement of paraspeckle components is required to maintain paraspeckle identity and prevent engulfment by other nuclear condensates^16^. Current models of paraspeckle assembly posit that NEAT1 is a triblock copolymer with distinct 5’/middle/3’ blocks^24^. Specific recruitment of NONO, and subsequent recruitment of FUS, to the middle of NEAT1 is thought to confer the middle region with relatively hydrophobic properties, causing NEAT1 to form layered micelles with the 5’/3’ ends in the shell and the middle region in the core^24^. However, the molecular preferences for paraspeckle proteins, including and beyond NONO, to bind and condense with distinct NEAT1 domains is poorly understood. More generally, dissecting and relating the multicomponent interaction logic amongst paraspeckle proteins and NEAT1 domains that endow paraspeckles with their emergent organization is a major open challenge.

In this work, we combine experiments, informatics, and physics-based simulations to examine how multiple proteins bind to different NEAT1 domains and assemble paraspeckle layers. Using fragments of NEAT1 from the shell-localized ends (5’/3’) and the core-localized middle region, we unexpectedly find that core proteins FUS and NONO preferentially bind and condense with the shell-localized 5’/3’ ends instead of the core-localized middle. The shell protein TDP-43 also binds the 3’ end preferentially, leading to a competition between core and shell proteins for end binding and weakening FUS and NONO condensation. Notably, TDP-43 competes less effectively with core proteins for the middle region, leading to surfactant-like TDP-43 layers on the surfaces of FUS/NONO condensates mirroring the ordering seen in paraspeckles. We develop a physical model that captures this interaction network and recapitulates layered organization through coarse-grained simulations. Our results prompt an update to the model of direct, selective binding between RBPs and their respective NEAT1 blocks. Instead, we propose that NEAT1 orders paraspeckle layers by directing the combinatorial recruitment of immiscible proteins that competitively bind different parts of NEAT1.

## Results

### Probing the current model of paraspeckle assembly

Previous work proposed that paraspeckle assembly is initiated when the core RBP NONO binds the assembly domain (8-16.6k) in the middle of NEAT1, specifically within three essential subdomains (9.8-12k, 12-13k, and 15.4-16.6k)^15,25^ (Fig. 1A). Subsequent recruitment of another core RBP, FUS, drives the adhesion of multiple NEAT1 transcripts through their middle domains to form core-shell paraspeckles (Fig. 1A). SFPQ also serves a key role in this process^15^ but ultimately ends up in both the core and shell^16^. We first asked if a simple physical description based on this model leads to paraspeckle-like assemblies. Towards this, we developed a coarse-grained molecular simulation with the following key features: (a) NEAT1 is modeled as a heteropolymer with three distinct monomer species that define the equal-sized 5’, middle, and 3’ blocks, (b) RBPs FUS and NONO are modeled as distinct species with spherical monomeric units, and (c) interactions between species i,j are modeled by short-range Lennard-Jones potentials with the parameter ɛ_*ij*_ that sets the strength of interactions (see Methods for details). We parameterized our simulation to reflect two essential but experimentally untested features of the current model: (i) NONO preferentially interacts with the middle region of NEAT1 compared to the ends, and (ii) FUS is recruited primarily by direct interactions with NONO rather than RNA^15^ (Fig. S1A; see Methods for simulation details). This coarse-grained representation, while foregoing molecular-scale sequence and interaction features, allows us to probe emergent mesoscale assemblies comprising thousands of molecules. Once parameterized, we ran Langevin dynamics with concentrations of biomolecules that loosely mimicked physiological, micromolar protein levels^26,27^ and nanomolar RNA levels, and averaged results over multiple trajectories (see Methods). As expected, we observed multi-layered clusters with NONO, FUS, and the middle regions of NEAT1 localized strongest to the core, while the 5’/3’ ends were more shell-localized (Fig. 1B and S1B). Fig. 1C shows averaged, normalized 2D projections of the cluster profiles for each component, highlighting this core-shell organization. Together, these data indicate that the assumptions underlying the current model are sufficient to generate paraspeckle-like, layered assemblies. However, direct experimental evidence that supports the above assumptions are lacking. We therefore set out to perform experiments using five, 1 kb fragments of human NEAT1: two from the 5’ and 3’ ends, named E1 and E2, respectively, and three from the essential subdomains^15^ of the middle region, named C1, C2, and C3 (Fig. 1D). We performed RNA binding and condensation assays with each RNA fragment using full-length, human paraspeckle proteins (Fig. 1D, see Methods).

**Fig. 1.**
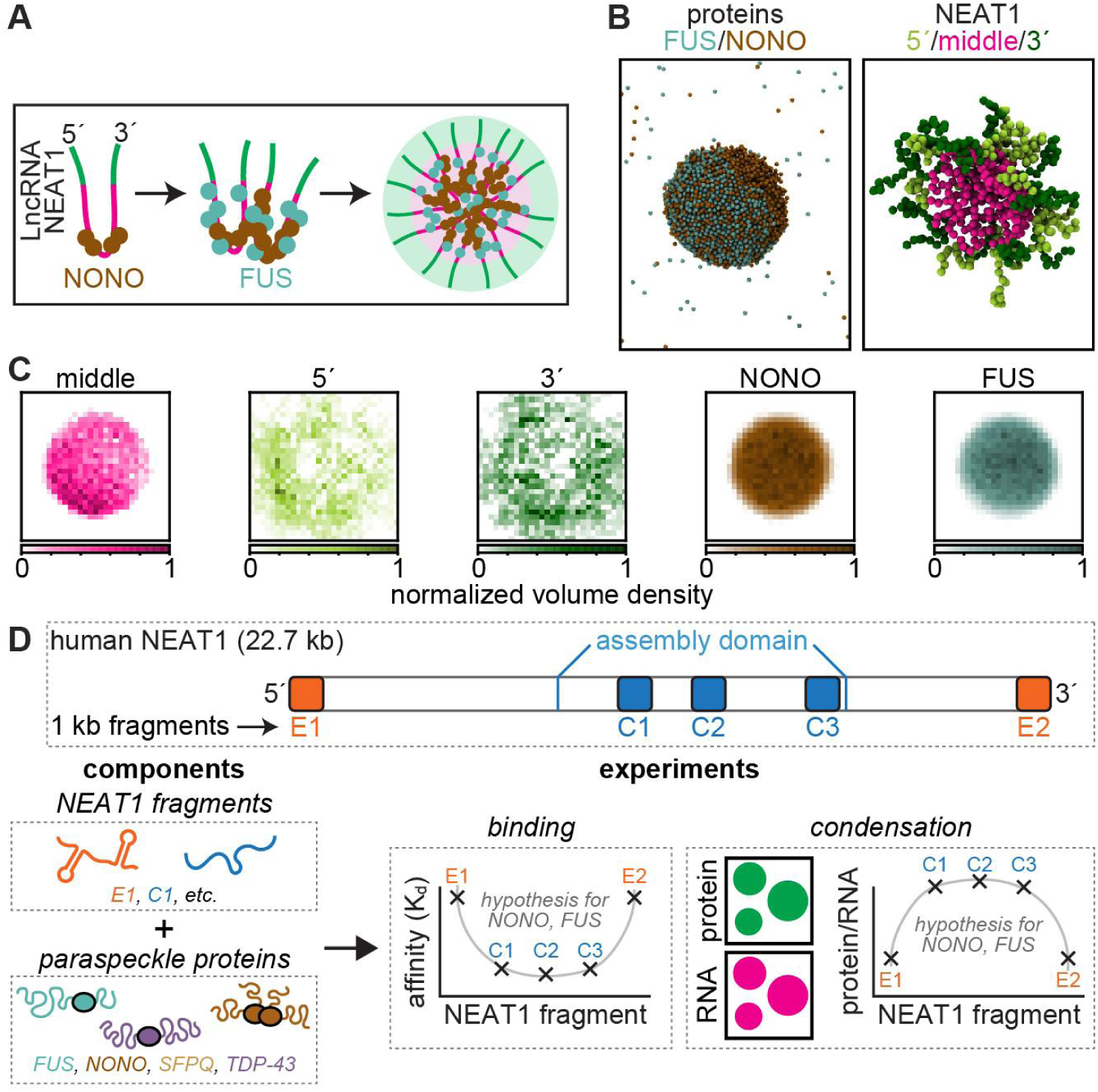
Probing the current model of paraspeckle assembly. (**A**) The current model suggests that the core proteins NONO and FUS localize selectively to the middle region of NEAT1, forming ribonucleoprotein units that adhere together to form paraspeckles. (**B**) Renderings of proteins (left) and NEAT1 chains (right) within the largest cluster at the end of a simulation. (**C**) Normalized maximum intensity projections (z-plane) of each component within clusters. Images represent average profiles of the largest clusters from five simulations. (**D**) Top: schematic of NEAT1 and the five *in vitro* transcribed 1 kb RNA fragments used in this study. Below: workflow of *in vitro* experiments. Plots depict hypothesized trends in RNA binding affinity and protein/RNA ratio within condensates for NONO and FUS.

### Paraspeckle core proteins exhibit unexpected binding preferences to NEAT1

We hypothesized that RNA binding motifs for NONO and FUS (Fig. 2A)^28,29^ should be more abundant in C1, C2, and C3 compared to E1 and E2. However, motif analysis of the NEAT1 sequence (Methods) did not reveal a clear enrichment for either protein within the assembly domain (Fig. 2A). Although a peak of moderate NONO motif density was apparent in the middle (Fig. 2A), the peak was located outside of the essential sub-domains required for layered paraspeckle assembly^15^. SFPQ also showed a lack of clear motif enrichment in the middle (Fig. 2A), consistent with its localization to both core and shell^16^. Unexpectedly, the regions of highest FUS motif density were at the NEAT1 ends, with the 5’ end showing the greatest density (Fig. 2A), contrasting with the known co-localization of FUS with the middle region in the core. Importantly, all of these observations appear consistent with eCLIP data from ENCODE^30^, in which NONO, SFPQ, and FUS exhibited binding peaks near the ends and a lack of obvious selectivity for the middle (Fig. S2A).

**Fig. 2.**
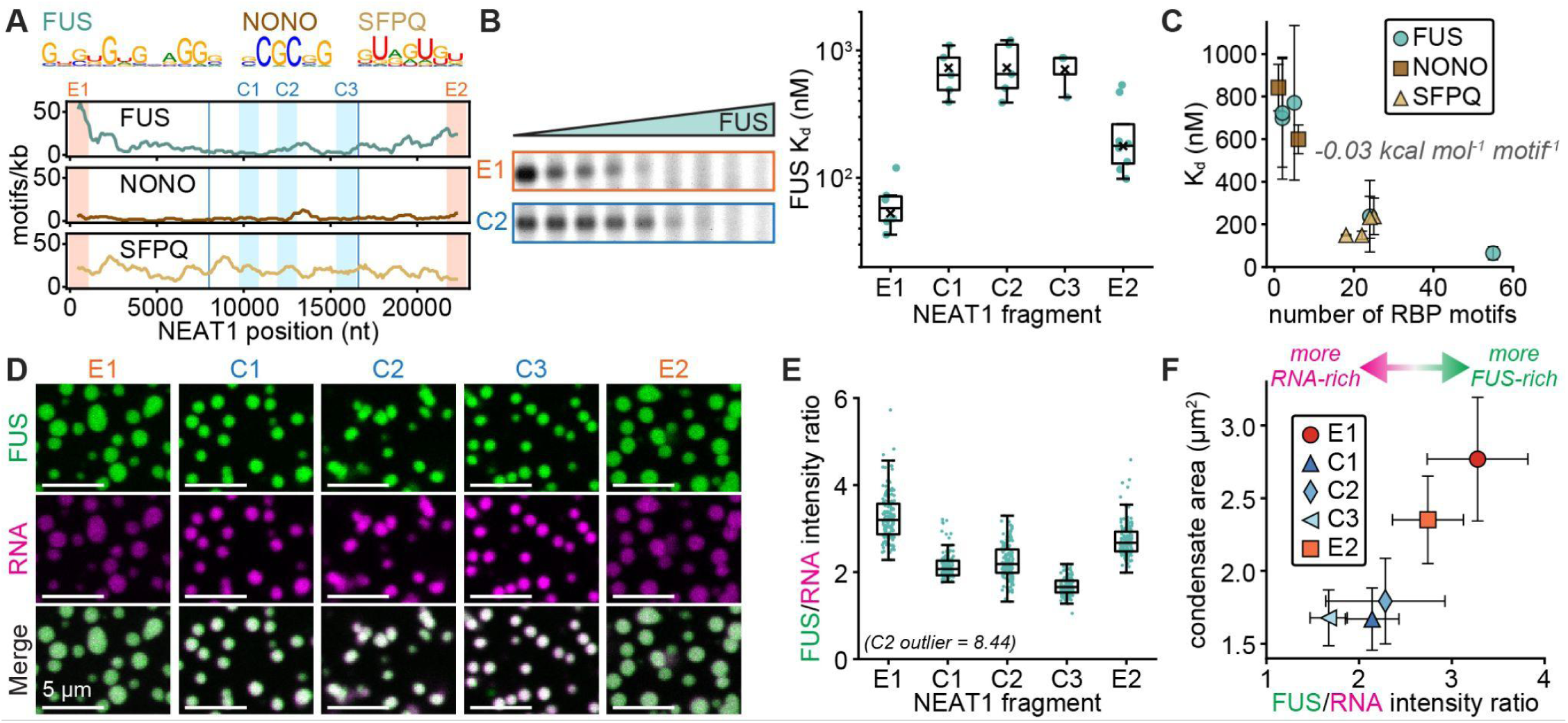
Paraspeckle core proteins exhibit unexpected binding and condensation with NEAT1. (**A**) FUS^29^, NONO^29^, and SFPQ^28^ RNA-binding motifs. Plots show the number of motifs for each protein within 1000 nt windows of NEAT1, tiled every 100 nt from 5’ to 3’. Orange and blue vertical bars indicate the five NEAT1 fragments used in this study, and the blue boxed region indicates the assembly domain identified in ref^15^ (8-16.6 kb). (**B**) Background-subtracted image of unbound RNA in FUS EMSA gel with E1 and C2 fragments. Plot: Apparent FUS-RNA dissociation constant (K_d_) estimates from EMSAs. Points and x’s represent fits to individual replicates and pooled data, respectively. (**C**) Apparent K_d_ as a function of the number of cognate RBP motifs per RNA. (**D**) Confocal slices of condensates assembled with 2 µM FUS-Atto 488 (green) and 30 nM RNA-Cy5 (magenta) after 3 h at 25 °C. FUS channel is contrasted identically in all images, while RNA channel is contrasted unequally to facilitate visual comparison of dense phase RNA concentrations after accounting for differences in RNA labeling density (Methods). (**E**) FUS/RNA intensity ratio at condensate centroids. Points represent individual droplets. (**F**) Condensate area as a function of FUS/RNA intensity ratio at condensate centroids. Data in (E, F) represent the 50 largest condensates from each experimental replicate (Methods). Boxes in (B, E) indicate interquartile range (IQR) with medians as bisecting lines and whiskers as 1.5*IQR. Points and error bars in (C, F) indicate mean ± one standard deviation.

To test if differences in motif abundance along NEAT1 impact protein binding, we performed gel electrophoresis mobility shift assays (EMSAs) with FUS^31^ and each NEAT1 fragment. These experiments revealed a generally greater affinity (i.e. lower apparent dissociation constant, K_d_) of FUS for the ends compared to the middle region (Fig. 2B). For example, FUS bound E1 with an order of magnitude greater affinity compared to C2, with apparent K_d_ of 65 ± 30 and 771 ± 363 nM, respectively (mean ± sd; Fig. 2B and S2B). Apparent RNA affinities of NONO and SFPQ were also generally consistent with cognate motif abundance, albeit with less distinctive differences between NEAT1 fragments compared to FUS (Fig. S2C-E). One notable exception was found with E2, which showed weak NONO binding despite containing the most NONO motifs of the tested RNAs (Fig. S2C). Thus, NONO binding is not explained by motif abundance alone, and likely relies on additional structural features^32^ that may not have been fully folded in EMSAs. Nonetheless, apparent K_d_ estimates for all proteins fell roughly along a common trend when plotted as a function of motif abundance (Fig. 2C), suggesting that each motif contributes similar, weak binding energy for its cognate protein. Recasting the data as ΔΔG of binding revealed a linear relationship with motif abundance, with a slope of –0.03 kcal mol^-1^ motif^-1^ (Fig. 2C and S2F). Thus, 20 motifs/kb strengthen binding by ∼0.6 kcal mol^-1^, or ∼1 k_B_T. The presence of more than 50 FUS motifs in E1 suggests that the 5’ end of NEAT1 is particularly tuned to bind FUS. However, these findings contrast with the current model of paraspeckle assembly that suggests a preference for the middle region. We asked if the binding differences among NEAT1 domains correlate with condensation preferences, or if distinct condensation behaviors may emerge that do not simply follow affinities.

### FUS condensates display NEAT1 sequence-dependent properties

To examine the NEAT1-dependent phase behavior of core proteins, we began by assembling condensates with 2 µM FUS (close to its approximate cellular concentration^26,27^) and each NEAT1 RNA fragment at varying concentrations (Fig. S3A). All RNAs promoted the formation of condensates (Fig. 2D and S3B), with FUS dense phase levels varying non-monotonically with RNA concentration (Fig. S3B, C), consistent with reentrant phase behavior^1^. Condensates became more rounded with smaller areas as RNA concentration increased (Fig. S3B, C), consistent with the formation of elastic RNA networks that may have slowed coarsening^33^. However, at RNA concentrations of 15, 30, and 60 nM, condensates assembled with middle region RNAs were consistently smaller compared to end RNAs (Fig. 2D and S3C, D), suggesting that middle region RNAs may have slowed coarsening more strongly. In line with this idea, RNA dense phase levels were consistently higher for middle region RNAs, while FUS levels were similar for all RNAs (Fig. S3E, F). As such, the FUS/RNA ratio (a proxy for the number of FUS molecules per RNA) was greater for the ends compared to the middle (Fig. 2E). The correlation between condensate area and FUS/RNA ratio (Fig. 2F) supports the conclusion that middle region RNAs formed denser RNA networks that slowed coarsening. In further support of this idea, secondary structure prediction of NEAT1 suggested that the middle region is more single-stranded (Fig. S3G) and therefore more capable of engaging in RNA-RNA interactions in the paraspeckle core^33,34^.

Condensate area scaled with the number of FUS motifs in each RNA (Fig. S4A), indicating that FUS has a previously unappreciated sequence affinity when condensing with RNA^35,36^. Based on this finding, we hypothesized that reducing the number of FUS motifs in an RNA should (i) weaken FUS binding and (ii) decrease condensate size. To test this, we compared wild-type E1 (E1^WT^, 55 FUS motifs) to a shuffled E1 sequence with fewer FUS motifs (E1^shuf^, 18 FUS motifs) (Fig. S4B). As expected, FUS bound E1^shuf^ with more than four-fold weaker affinity compared to E1^WT^ (Fig. S4C), and FUS condensates assembled with E1^shuf^ were smaller compared to E1^WT^ (Fig. S4D). Thus, FUS condensate area scaled with apparent RNA affinity for all tested RNAs (Fig. S4E). Together, these findings reveal that FUS displays NEAT1 sequence-dependent phase behavior without a clear preference for the middle, forming larger, more protein-rich condensates in the presence of NEAT1 regions with stronger FUS binding. Given that the current model of paraspeckle assembly suggests a sequential recruitment of NONO to the middle, followed by FUS (Fig. 1A)^15^, we wondered if NONO may condense preferentially with the NEAT1 middle region.

### The NEAT1 ends promote NONO condensation

To examine if NONO condensation is NEAT1 sequence-dependent, we repeated condensate experiments with purified NONO. In the absence of RNA, NONO formed spherical condensates at a concentration of 2 µM (Fig. 3A), indicating that NONO engages in homotypic interactions that drive protein-only condensation, similar to the related proteins SFPQ and PSPC1^37,38^. Addition of E1 or E2 RNAs stimulated NONO condensation, increasing condensate size and NONO dense phase levels (Fig. 3A, B and S5A). Strikingly, E1 and E2 RNAs also formed RNA-rich sub-phases localized near the condensate periphery (Fig. 3A), suggesting that NONO affinity for E1 and E2 was sufficient to drive co-condensation initially but insufficient to maintain a well-mixed state, as observed for other protein-RNA condensates^39^. Curiously, NONO did not co-condense with middle region RNAs, instead forming protein-rich but RNA-depleted condensates that were trapped within amorphous RNA networks (Fig. 3A). This finding strongly contrasts with the current model of paraspeckle assembly (Fig. 1A), leading us to wonder if NONO requires additional, cooperative interactions with FUS in order to localize to the paraspeckle core.

**Fig. 3.**
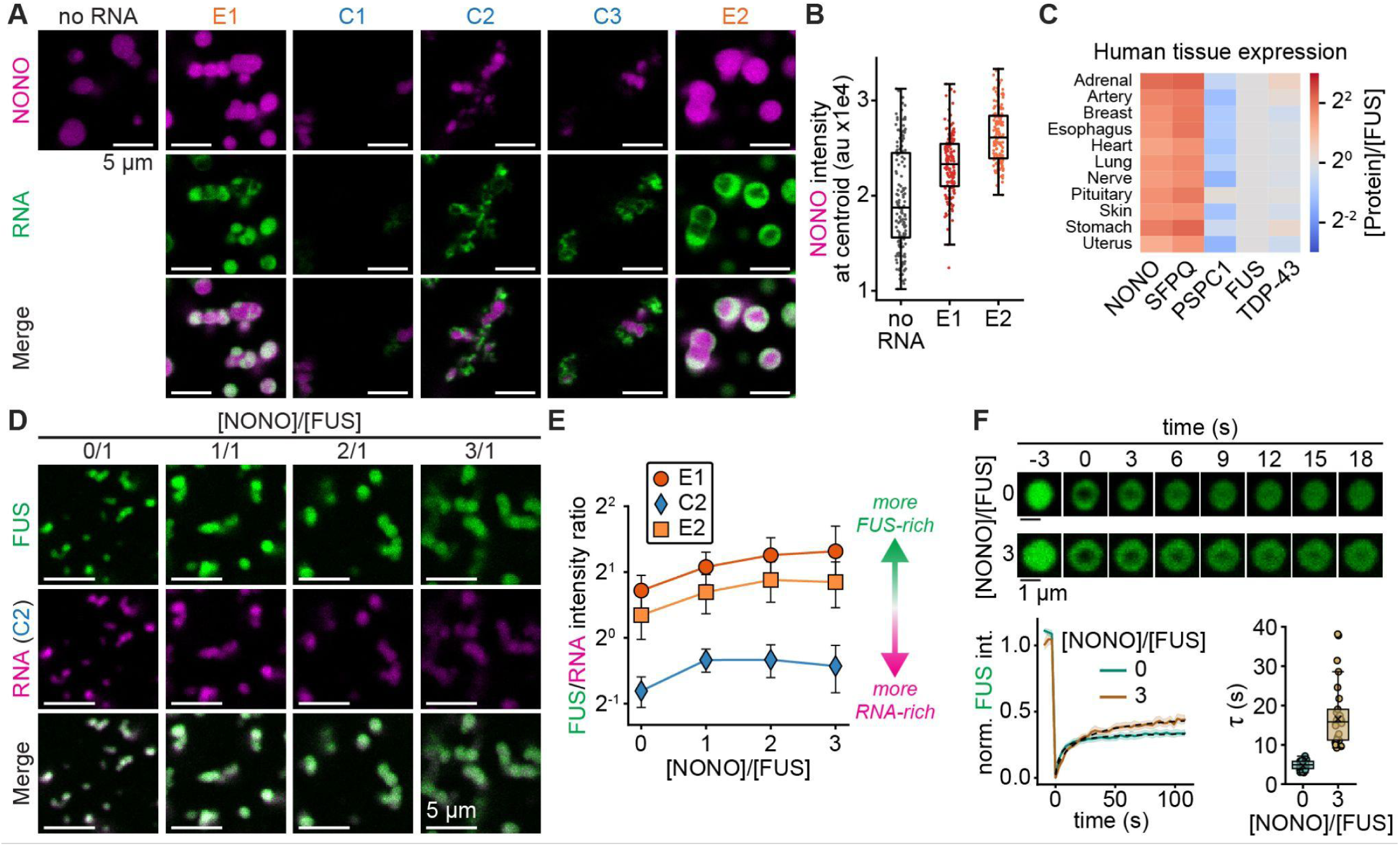
NONO/FUS stoichiometry modulates condensation. (**A**) Confocal slices of condensates assembled with 2 µM NONO (magenta) and 15 nM RNA (green) after 3 h at 25 °C. NONO and RNA channels are contrasted equally in all images. (**B**) NONO intensity at condensate centroids from experiments corresponding to (A). Data represent the 50 largest condensates from each experimental replicate (Methods). Points indicate individual condensates, boxes indicate IQR with medians as bisecting lines and whiskers as 1.5*IQR. Core RNAs were not included in analysis owing to variable z positions of NONO condensates trapped within RNA networks. (**C**) Heat map of the abundance of the indicated paraspeckle proteins relative to FUS in human tissues, quantified using data from ref^41^. (**D**) Confocal slices of condensates assembled with 1 µM FUS (green) + 15 nM C2 RNA (magenta) + NONO at the indicated concentrations after 3 h at 25 °C. FUS and RNA channels are contrasted equally in all images. (**E**) FUS/RNA intensity ratio at condensate centroids as a function of NONO/FUS stoichiometry with the indicated RNAs. Data points and error bars indicate mean ± one standard deviation for the 50 largest condensates from each experimental replicate. (**F**) Top: confocal slices of FUS (green) photobleaching recovery within droplets assembled with 1 µM FUS + 15 nM E2 RNA + NONO at the indicated stoichiometry. Lower left: normalized FUS intensity recovery profiles as a function of time for droplets with the indicated NONO stoichiometry. Curves and shaded error bars indicate average ± 95% CI, dashed lines indicate fits to single component FRAP recovery model^44^. Lower right: FUS recovery time constant (τ) estimates from FRAP fits. Circles and x’s indicate τ from fits to individual droplets and average data, respectively. Boxes indicate IQR with medians as bisecting lines and whiskers as 1.5*IQR.

### A multicomponent stoichiometry modulates condensation

Since stoichiometry is a key modulator of condensate behavior^40^, we first sought to understand the relative concentrations of FUS and NONO in cells. Analysis of paraspeckle protein abundance from published, quantitative proteomics data^41^ revealed that NONO and the related protein SFPQ were consistently more abundant than FUS in most human tissues by a factor of 2-3 (Fig. 3C). Data from different cell lines^42^ and mammalian species^43^ yielded similar results (Fig. S5B), indicating that this NONO/FUS stoichiometry is a shared feature of many cell types across species. When mixed at a physiological stoichiometry of 3/1, NONO and FUS co-partitioned into condensates with both end and middle region RNAs, with similar dense phase levels of each protein across the different RNAs (Fig. S5C). Thus, FUS-NONO interactions at physiological stoichiometries can overcome the tendency of NONO to demix from the middle region.

To understand how NONO modulates FUS-NEAT1 phase behavior, we varied NONO concentration while keeping FUS and RNA concentrations fixed. Fig. 3D shows condensates with C2 RNA, where inclusion of NONO led to progressively larger condensates with lower FUS and RNA levels (Fig. S5E). Similar results were obtained with E1 and E2 RNAs (Fig. S5D, E). However, the reduction in RNA appeared to outpace that of FUS for all RNAs, thereby increasing FUS/RNA ratio as NONO was added (Fig. 3E). This finding suggests that although FUS and RNA were diluted and/or evicted to accommodate NONO, RNA was evicted to a greater extent owing to relatively weak NONO-RNA binding. In line with this idea, NONO did not substantially impact FUS-RNA binding (Fig. S5F). Thus, NONO likely interacts more strongly with FUS compared to RNA. These NONO-FUS interactions appeared to reduce FUS mobility, as evidenced by a more than three-fold reduction in the FRAP recovery time constant of FUS (5 ± 1 and 18 ± 8 s without and with NONO, respectively, mean ± sd; Fig. 3F). Interestingly, the FUS mobile fraction increased approximately 1.5-fold, from 0.35 ± 0.07 to 0.51 ± 0.12 (mean ± sd; Fig. S5G), and NONO displayed poor recovery overall (Fig. S5H). Together, these data suggest that NONO forms a relatively stable network in condensates that lowers FUS mobility but increases the pool of mobile FUS. However, the similar dense phase levels of FUS and NONO with the NEAT1 ends and middle (Fig. S5C) suggests that NONO-FUS interactions do not impart selectivity for the middle. We next sought to understand what might prevent NONO and FUS from accumulating in the paraspeckle shell.

### The shell protein TDP-43 opposes FUS and NONO condensation by competing for RNA binding

As TDP-43 is the canonical paraspeckle shell protein^14^, we hypothesized that TDP-43 might help to exclude FUS and NONO from the NEAT1 ends in the shell. TDP-43 recruitment to paraspeckles is driven by at least three UG repeat regions in NEAT1^45^. As expected, TDP-43 motif^29^ analysis of NEAT1 revealed several peaks centered near these UG repeats which aligned with eCLIP binding peaks from ENCODE (Fig. 4A, S6A). The most prominent peak was in a shell-localized region^13,24^ near the 3’ end (21.1-22.1k, 0.6k upstream of E2). We named this 3’ end-proximal, 1 kb RNA fragment E3 (Fig. 4A). To confirm that TDP-43 binds E3, we *in vitro* transcribed wild-type E3 (E3^WT^, 56 TDP-43 motifs) and a shuffled version with fewer motifs (E3^shuf^, 5 TDP-43 motifs) (Fig. S6B). TDP-43^46^ bound E3^WT^ with apparent K_d_ of 150 ± 78 nM (mean ± sd), but did not bind E3^shuf^ with measurable affinity (Fig. S6B). Weak TDP-43 binding to E1 and C2 also hindered K_d_ estimation for these RNAs (Fig. S6C). Thus, TDP-43 binds selectively to UG repeat motifs in NEAT1 E3.

**Fig. 4.**
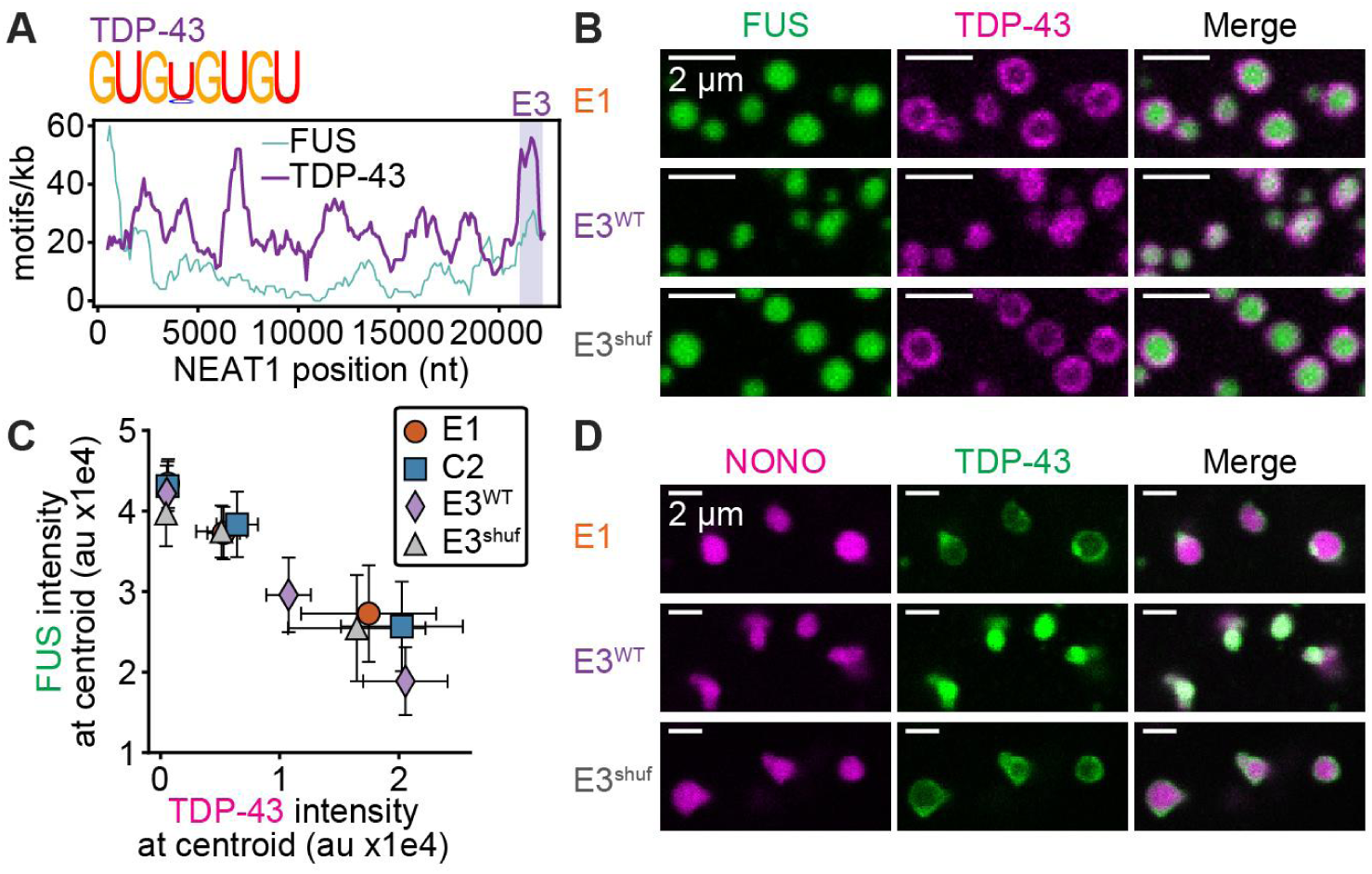
TDP-43 opposes core protein condensation by competing for RNA binding. (**A**) TDP-43 RNA binding motif^29^ and plot of the number of FUS and TDP-43 motifs within 1000 nt windows of NEAT1, tiled every 100 nt. FUS motif distribution repeated from Fig. 2A. Purple vertical bar indicates E3 fragment. (**B**) Confocal slices of condensates assembled with 2 µM FUS (green) + 0.2 µM TDP-43 (magenta) + 15 nM of the indicated, unlabeled RNA after 3 h at 25 °C. (**C**) FUS intensity as a function of TDP-43 intensity at condensate centroids. Data are from experiments in which TDP-43 was added at concentrations of 0, 0.2, and 0.5 µM, with fixed FUS and RNA concentrations indicated in (B). Data points and error bars indicate mean ± one standard deviation for the 50 largest condensates from each experimental replicate (Methods). (**D**) Confocal slices of condensates assembled with 2 µM NONO (magenta) + 0.2 µM TDP-43 (green) + 15 nM of the indicated, unlabeled RNA after 3 h at 25 °C.

Of note, FUS bound E3^WT^ with similar affinity to TDP-43 (Fig. S6D), suggesting that the two proteins may compete for binding to the shell-localized NEAT1 3’ end. Based on a recent report that TDP-43 can disperse FUS condensates^47^, we hypothesized that such a competition may inhibit FUS condensation. To test this, we coassembled condensates with FUS, TDP-43, and several NEAT1 fragments (Fig. 4B). Strikingly, TDP-43 was excluded from condensates assembled with most RNAs, forming TDP-43-rich shells surrounding FUS-rich cores, reminiscent of paraspeckles (Fig. 4B and S6E). However, TDP-43 was recruited into FUS condensates assembled with E3^WT^ (Fig. 4B and S6F), suggesting that strong RNA binding can overcome the tendency of FUS to exclude TDP-43. Indeed, TDP-43 dense phase levels scaled with the number of TDP-43 motifs in each RNA (Fig. S6G), indicating that TDP-43 recruitment into condensates depends on cognate motif abundance. Importantly, TDP-43 recruitment weakened FUS condensation, as condensate area and dense phase FUS levels both scaled inversely with TDP-43 levels for all RNAs (Fig. 4C and S6H). We obtained similar results when adding TDP-43 to pre-assembled FUS-NEAT1 condensates (Fig. S7A-C), in which TDP-43 was recruited strongest into E3^WT^ condensates and reduced FUS levels over time (Fig. S7D-F). Finally, TDP-43 formed similar, surfactant-like layers on the surfaces of NONO-NEAT1 condensates (Fig. 4D), reducing condensate size and NONO levels (Fig. S8). Thus, immiscibility with TDP-43 is a shared feature of paraspeckle core proteins. These data support the hypothesis that competitive TDP-43 binding to the NEAT1 3’ end creates an unfavorable environment that may exclude FUS and NONO from the shell. To further test this hypothesis, we returned to our coarse-grained simulations of paraspeckle assembly.

### An updated model relates multicomponent biomolecular interactions to paraspeckle assembly

To relate the multicomponent interactions observed in our experiments to paraspeckle assembly, we again used our coarse-grained physical model, updated here to include TDP-43 as a distinct species. Based on experimental observations (Figs. 2-4), we modified the intermolecular interactions to reflect (a) NEAT1 end binding by all RBPs, (b) weak protein-protein interactions between FUS/NONO and TDP43, and (c) stronger RNA-RNA interactions between NEAT1 middle regions, reflecting its predicted single-stranded nature (Fig. 5A and S9A, Methods). Finally, we updated the concentrations of molecules to roughly reflect the estimated excess of NONO relative to FUS, and we ran Langevin dynamics. Simulations showed the formation of multi-layered clusters with NONO/FUS in the core and TDP-43 in the shell (Fig. 5A, B and S9B). Of note, TDP-43 formed discrete, patch-like clusters in the shell (Fig. 5A, B), likely to minimize middle region interactions. Interestingly, TDP-43 appeared similarly patchy in previous imaging studies^14^, suggesting that our simulations may reflect physiologically-relevant TDP-43 localization. However, such TDP-43 patches did not appear to bind the ends of every NEAT1 chain, leaving some end segments free to bind NONO/FUS in the core. As a result, NEAT1 domains were less clearly delineated into core and shell layers compared to proteins (Fig. 5B), though the middle region was more core-enriched compared to the ends (Fig. S9B). Taken together, these findings support the hypothesis that although both core and shell proteins bind preferentially to the NEAT1 ends, TDP-43 immiscibility with NONO/FUS establishes a competition that inhibits NONO/FUS condensation with the ends (Fig. 5C). Thus, we propose that the experimentally-motivated interaction network of our updated model can give rise to paraspeckle-like, layered assemblies.

**Fig. 5.**
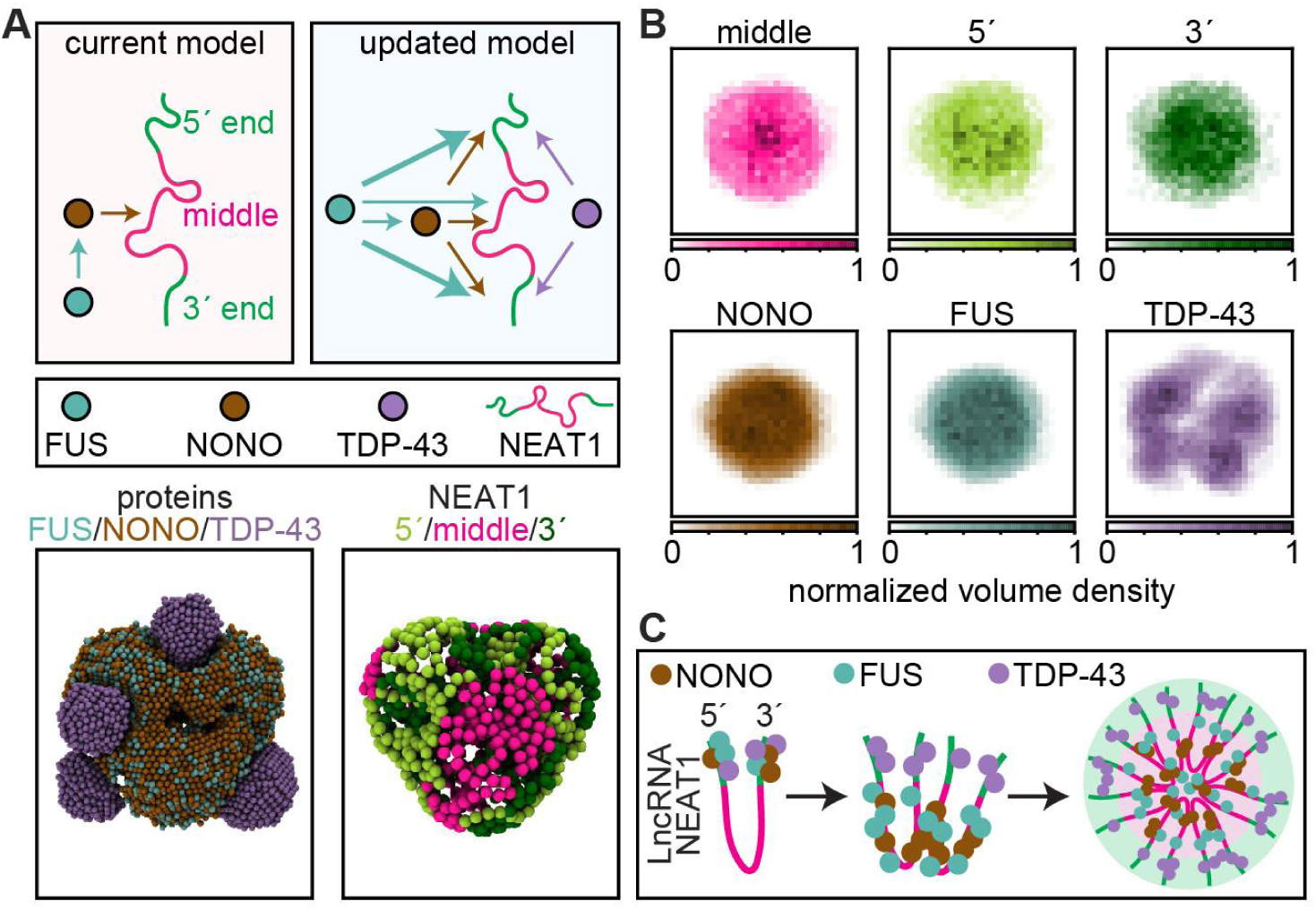
An updated model of paraspeckle assembly. (**A**) Summary of key interactions in the updated model. Thicker arrows correspond to stronger interactions. Below: renderings show proteins (left) and NEAT1 chains (right) within the largest cluster at the end of a simulation. (**B**) Normalized maximum intensity projections (z-plane) of each component within clusters. Images represent average profiles of the largest clusters from five simulations. (**C**) Conceptual schematic of the updated model.

As a final note, since paraspeckles form co-transcriptionally^48^, we asked if active transcription might be important for guiding assembly. To emulate transcription, we simplified our model to an RNA-only description with enhanced middle region interactions. Active transcription was modeled through adding new chains as the simulation progressed, and the relative rate of transcription to chromatin motion was mimicked by modulating the genomic region in which subsequent 5’ ends were colocalized (see Methods for details). We found that the transcriptional dynamics of NEAT1 strongly determine the number of RNA chains incorporated into clusters in our simulations (Fig. S10). Specifically, a transcriptional burst, in which multiple NEAT1 copies accumulate in close spatial proximity, led to multi-RNA clusters reminiscent of paraspeckles, whereas slow transcription led to small clusters containing fewer chains (Fig. S10). Thus, we emphasize that the biochemical interactions described above likely take place in the context of co-transcriptional assembly, with RBPs binding nascent NEAT1 RNAs as they are synthesized in a transcriptional burst.

## Discussion

Diverse condensates display internal organization into spatial subdomains with distinct molecular identities. Here we uncovered a network of interactions between multiple proteins and the NEAT1 lncRNA that can give rise to paraspeckle layers. Unexpectedly, the essential core proteins FUS and NONO display binding and condensation preferences with the shell-localized NEAT1 ends (Fig. 2 and 3), suggesting that additional factors are needed to prevent core proteins from localizing to the shell. We propose that the shell protein TDP-43 serves such a function by competing with FUS and NONO for the ends and weakening core protein condensation. TDP-43 competes less effectively with FUS and NONO for the NEAT1 middle region, leading to TDP-43 shells surrounding FUS and NONO cores (Fig. 4). Thus, we propose that competitive RNA binding between immiscible proteins leads to core-shell arrangement as seen in cells (Fig. 5C).

The molecular mechanisms of multi-domain condensation are a topic of ongoing investigation. One of the most well-studied multi-domain condensates is the nucleolus, in which differences in subdomain properties such as surface tension and hydrophobicity drive their spatial ordering^6,49^. Such properties emerge from distinct protein sequence features associated with each subdomain. We hypothesize that FUS/NONO and TDP-43 also possess distinct sequence characteristics that encode different emergent condensate properties which remain to be carefully dissected.

The RNA binding specificity of FUS is debated, with some reports suggesting no sequence preferences and others suggesting specificity for some sequence and structural elements^35,50,51^. Here we find that low-affinity, degenerate sequence motifs can strengthen FUS binding in proportion to their local density. Although individual motifs do not substantially impact binding, we propose that local regions of high motif density in NEAT1 may be tuned to strongly bind FUS and other paraspeckle proteins (Fig. 2A). Such additive effects of multiple, low-affinity motifs resemble transcription factor binding at DNA regulatory elements, in which repetitive, low-affinity regions of DNA can extend transcription factor dwell time and assist the search for high-affinity consensus motifs^52^. Thus, we propose that NEAT1 contains an embedded motif grammar that can dictate combinatorial RBP binding to assemble layers.

In addition to the protein-RNA interactions encoded by NEAT1, we propose that sequence-encoded RNA-RNA interactions among NEAT1 chains are also key for paraspeckle assembly. Specifically, our results suggest that the middle region of NEAT1 is more single-stranded compared to the ends, and therefore more capable of engaging in RNA-RNA interactions that might help establish a distinct material state in the core. Thus, NEAT1 may contain an additional RNA-RNA interaction grammar that contributes to paraspeckle layering.

Finally, paraspeckle assembly is a co-transcriptional process in which RBPs bind nascent NEAT1 transcripts as they are synthesized^48^. We propose that transcriptional bursts are required to form paraspeckle-like clusters containing many copies of NEAT1 (Fig. S10). Paraspeckle abundance and morphology can change in response to a variety of stimuli including stress and infection^53^, suggesting that such stimuli may alter transcriptional burst size and/or frequency. The relationship between NEAT1 transcriptional burst parameters and paraspeckle morphology and subphase organization may be a key avenue of future study.

## Acknowledgements

A.S.G. acknowledges support from the Air Force Office of Scientific Research (FA9550-20-1-0241), the NIH (R35GM156800 and R01GM081506), and the NSF (MCB 2044895). W.T.S. acknowledges a Pathway to Independence Award from NIH (K99GM149757). M.K.S. was supported in part by the NIH Training Grant (T32GM153505) through Northwestern University’s Biotechnology Training Program. K.S. acknowledges support from Northwestern for startup funding. M.K.S. and K.S. acknowledge support in part through the computational resources and staff contributions provided for the Quest high performance computing facility at Northwestern University.

## Contributions

W.T.S. and A.S.G. designed *in vitro* binding and condensation experiments. W.T.S. performed all experiments and analyzed experimental data. W.T.S and K.S. conducted all informatics analyses reported. M.K.S. and K.S. designed and performed simulations of paraspeckle assembly and analyzed simulation data. All authors wrote and edited the manuscript.

## Data and code availability

Data and code that accompany this manuscript are available at: https://github.com/shrinivaslab/2025_snead_paraspeckles Raw, unanalyzed images and other data files supporting the findings of this study are available from the corresponding authors on reasonable request.

## Methods

### Computational model of paraspeckle assembly

#### Coarse-grained molecular dynamics simulations

We employ coarse-grained molecular dynamics simulations^54^ to model paraspeckle formation. In our model, we model the following paraspeckle components as distinct species: (a) long noncoding RNA NEAT1, (b) DBHS family proteins SFPQ, NONO, and PSPC1 are modeled as a single species, and to motivate connections to experiments, labeled as NONO unless otherwise specified, (c) FUS protein, and (d) TDP-43 protein. NEAT1 is modeled as a heteropolymer (N=45 units) where each monomer on the chain represents approximately 500 bp and has a diameter of 1 unit. Sequence features of NEAT1 are captured in monomer identity. Unless otherwise specified, NEAT1 is modeled as a polymer with 3 distinct monomers that comprise equi-sized domains (l=15 units) in the 5’, middle, and 3’ regions. The various proteins are represented as isotropic monomers with a diameter of ½ unit. This coarse-grained representation enables us to elucidate paraspeckle assemblies with thousands of constituents.

Effective interactions between particles are modeled using a shifted Lennard Jones potential utilizing the Hoomd-Blue LJ module^54^. Bonded interactions between monomers of NEAT1 are modeled as a harmonic potential with a spring constant of k=500. Energy units in the simulation are scaled to a kT of 1. The initial conformation of NEAT1 chains in simulations are initialized as a self-avoiding random walk and the various protein species are then randomly positioned in the simulation box. A typical simulation is run with 20 chains of NEAT1 (*c*_*RNA*_ ≈ 10*nM*) and 12000 copies (*c*_*protein*_ ≈ 6µ*M*) of total protein with equal concentration of each species unless otherwise specified to loosely mimic physiological scales.

Individual trajectories are run with Langevin dynamics (γ = 1) and periodic boundary conditions for 2.5 × 10^8^ steps with *dt* = 8 × 10^-3^. Once trajectories start the last 6. 25 × 10^7^ steps, snapshots are logged every 6. 25 × 10^5^ steps in a GSD file. The last 75 frames from these snapshots are used for statistical analysis of cluster properties discussed below and data reported are averaged over multiple trajectories (n=5). All simulations were performed on GPUs (40 GB PCIe and 80 GB SXM cards) from the Northwestern Quest High-Performance Computing Cluster.

#### Coarse-grained simulations of co-transcriptional assembly

To study co-transcriptional assembly of paraspeckles (Fig. S10), we consider an RNA-only model with effective interactions between middle-regions of NEAT1 driving assembly. To mimic active and burst transcription, individual chains were added to the simulation at 1 copy per a specified number of steps. Once 50 copies was reached, no further copies were added until the end of simulation (5 × 10^5^ steps). Fast bursts of transcription were modeled by initiating 5’ ends of newly transcribed NEAT1 every 100 steps within a small spatial neighborhood of size (6. 25 *units*^3^ ≈ 65 × 65 × 65 *nm*^3^). Slow bursts of transcription were modeled by initiating 5’ ends of newly transcribed NEAT1 every 800 steps within a large spatial neighborhood (of size (150 *units*)^3^ ≈ 1. 5 × 1. 5 × 1. 5 µ*m*^3^). In this slower model, the tacit assumption is that transcription is much slower than rate of chromatin motion, explicitly modeled by both a slower rate of transcription and faster motion of active gene loci.

#### Interaction network parameters

The choice of LJ potential well-depth is used to capture an effective interaction between multiple species – represented in matrix form (Fig. S1A and S9A) and parameter values are provided (Table S1).

##### Current Model

To represent the current model of paraspeckle assembly (Fig. 1), we included interactions between NONO and the middle region of NEAT1 (M) and between FUS and NONO with minimal interactions otherwise (Fig. S1A).

##### Updated Model

Based on experimental findings reported in this paper (Fig. 2-4), an updated model was developed (Fig. 5 and S9). Key components of this interaction matrix include the following:

● Strong RNA-RNA interactions between the middle regions of NEAT1 (ɛ_*M*−*M*_ = 2*kT*), as suggested by Fig. 2E, F and S3G, which suggest high single-stranded content in the middle regions.
● NONO has relatively non-specific affinity for the entire length of NEAT1 as suggested by Fig. S2C.
● FUS has an asymmetrical affinity across NEAT1 with affinity for the 5’ end, 3’ end, and middle of the polymer in decreasing order respectively (Fig. 2B).
● Synergistic NONO and FUS interactions, as reported in Fig. S5C.
● TDP-43 only has high self-interactions and specific affinity for the 5’ and 3’ end of NEAT1 as shown in Fig. 4A and S6B, C. While our experiments revealed strong TDP-43 binding only to E3 fragments (in 3’ end), eCLIP data (Fig. S6A) indicates that TDP43 is likely interacting with both 5’ and 3’ ends.
● We report simulations with a higher concentration of NONO/FUS (2/1).

#### Cluster profiles

Analysis of the data was mainly done as post-simulation calculations except for ones employed in Hoomd-Blue during simulation including logs of potential energy, kinetic energy, and temperature^54^. Post-simulation calculations were performed using a combination of python packages including *numpy*, *freud*, and *matplotlib*. Data was stored in GSD files and read through the gsd module on python. Generally analysis included determining clusters with a mix of protein and RNA together using the freud cluster module under a neighboring threshold of 1.5 unit distance^55^. For each cluster determined, properties including the center of mass, size and radius of gyration (rg) were extracted, and used for the specific analysis explained below. Each cluster property including radial density plots (RDP) and maximum intensity plots (MIP) are averaged over the largest clusters in each trajectory, cumulating from all specified frames, then averaged over multiple runs.

#### Radial density plots

For each frame used for analysis, all specified clusters with protein and RNA are reported with a rg and center of mass. From the center of mass to 2*rg of clusters, all particle’s distance from the center for a specific species (5’, M, 3’, NONO, FUS, and TDP-43) were organized into bins (number of bins=10). These bins are then divided by its representative spherical volume to give a number density. The radial density for each species is plotted over the normalized distance from the center of the cluster. This was visualized using matplotlib in Python.

#### Maximum intensity plots

For each frame used for analysis all specified clusters with protein and RNA are reported with its rg and center of mass. From the center to 2*rg of clusters, the plane of either XY, YZ, or ZX is divided into a 30×30 grid. For a plane’s orthogonal axis it is divided into three hundred slices. For each slice, the 30×30 grid is filled with a specific species local density that is calculated from freud’s local density module (r_max=1 and diameter=1)^55^. Over all slices, each cell in the grid records its largest density. The maximum intensity plots for each species is visualized using matplotlib heat maps and normalized to each species highest recorded density.

#### Visualization of simulation data

Figures were generated using Visual Molecular Dynamics (VMD)^56^. Images were rendered using Tachyon (Internal) with RNA monomers as AOChalky and proteins as AOShiny into tga files. The python package, *pillow*, was used to convert the tga files into pdfs.

##### Chemical materials

HEPES, LiCl, DEPC, TCEP, bovine serum albumin (BSA), β-mercaptoethanol, Pluronic-F127, glycerol, imidazole, ATP disodium salt hydrate, Atto 488-NHS-ester, Atto 594-NHS-ester, and Cy5-UTP were purchased from Sigma-Aldrich. NaCl, KCl, EDTA, Triton X-100, L-arginine, and Pierce EDTA-free protease inhibitor tablets were purchased from Thermo Fisher Scientific. IPTG was purchased from Gold Biotechnology. MgCl_2_ was purchased from VWR. Atto 488-UTP was purchased from Jena Bioscience.

##### RBP motif analysis of NEAT1

RNA-binding protein (RBP) motif analysis of RNA sequences was performed using the Bio.motifs package from biopython version 1.85 (https://biopython.org/docs/dev/api/Bio.motifs.html). Position-weight matrix (PWM) representations of RBP motifs were created by entering “counts” for each nt at each motif position. For FUS, NONO, and TDP-43, counts were obtained from ref^29^ For SFPQ, counts were obtained manually from the sequence logo provided by ref^28^ (https://hugheslab.ccbr.utoronto.ca/supplementary-data/RNAcompete_eukarya/Experiment_reports/RNAcompete_report_index.html). Briefly, nucleotide (nt) heights at each position were measured using ImageJ. The nt heights in pixels served as proxies for relative nt counts. Sequence logos for each PWM were created using the logomaker package in Python. The reconstructed SFPQ logo closely matched the motif presented in ref^28^.

PWMs were converted to position-specific scoring matrices (PSSMs), which were used to search for motifs in human NEAT1 (NCBI Accession NR_131012) with false-positive rate set to 1%. The number of motifs within 1000 nt windows of NEAT1, tiled every 100 nt, were summed to create plots of RBP motif density.

##### Secondary structure analysis of NEAT1

RNA secondary structure prediction was performed using the ViennaRNA package for Python version 2.7.0 (https://www.tbi.univie.ac.at/RNA/ViennaRNA/doc/html/index.html). 1000 nt fragments of NEAT1, tiled every 100 nt, were folded and the minimum free energy (MFE) of the optimal secondary structure was obtained and plotted. To estimate pair probability of each NEAT1 fragment, matrices of base pair probabilities were obtained and summed along the rows to obtain the total probability of base pairing for each nucleotide in the fragment. The average ± standard error of the mean of the resulting list of pair probabilities was used for plotting.

##### Analysis of quantitative proteomics data

Previously published quantitative proteomics datasets were obtained for human tissues^41^, multiple cultured cell lines^42^, and matched orthologs in primary skin fibroblasts from different mammalian species^43^. Briefly, from each of these data, protein counts were averaged across replicates and normalized to count of FUS. These effective expression ratios are reported in Fig. 3C and S5B.

##### Plasmids

Plasmids harboring NEAT1 fragment templates for *in vitro* transcription were ordered from Twist Bioscience (E1^WT^, E1^shuf^, C1, C2, C3, and E2) in the pTwist Amp High Copy vector, or GenScript (E3^WT^ and E3^shuf^) in the pUC57 vector. All 1000 nt sequences contained identical 5’/3’ handles (5’-GAGTTCTACAGTCCGACGATc-3’ / 5’-tGGAATTCTCGGGTGTCAAGG-3’) for PCR amplification of templates.

Bacterial expression plasmids encoding 6xHis-MBP-FUS-FL-WT (FUS) and TDP-43-FL-WT-MBP-6xHis (TDP-43) were gifts from N. Fawzi (Addgene no. 98651 and 104480). The bacterial expression plasmid encoding MBP-7xHis-TEV protease (TEV) was a gift from D. Waugh (Addgene no. 8827). Bacterial expression plasmids encoding 6xHis-MBP-NONO FL and 6xHis-MBP-SFPQ FL (NONO and SFPQ) were cloned using the FUS vector. Specifically, the FUS insert was excised using NdeI/XhoI enzymes, and NONO and SFPQ inserts containing 5’/3’ NdeI/XhoI sites were PCR amplified from pcDNA3.1 vectors harboring each gene (Addgene no. 127655 and 166959, respectively, gifts from N. Manel and R. Segal). Following restriction digestion with NdeI/XhoI enzymes, NONO and SFPQ inserts were ligated into the linearized vector using T4 DNA ligase (NEB). All plasmid constructs were confirmed using Sanger and/or whole plasmid sequencing (Genewiz). Primer sequences for cloning were:

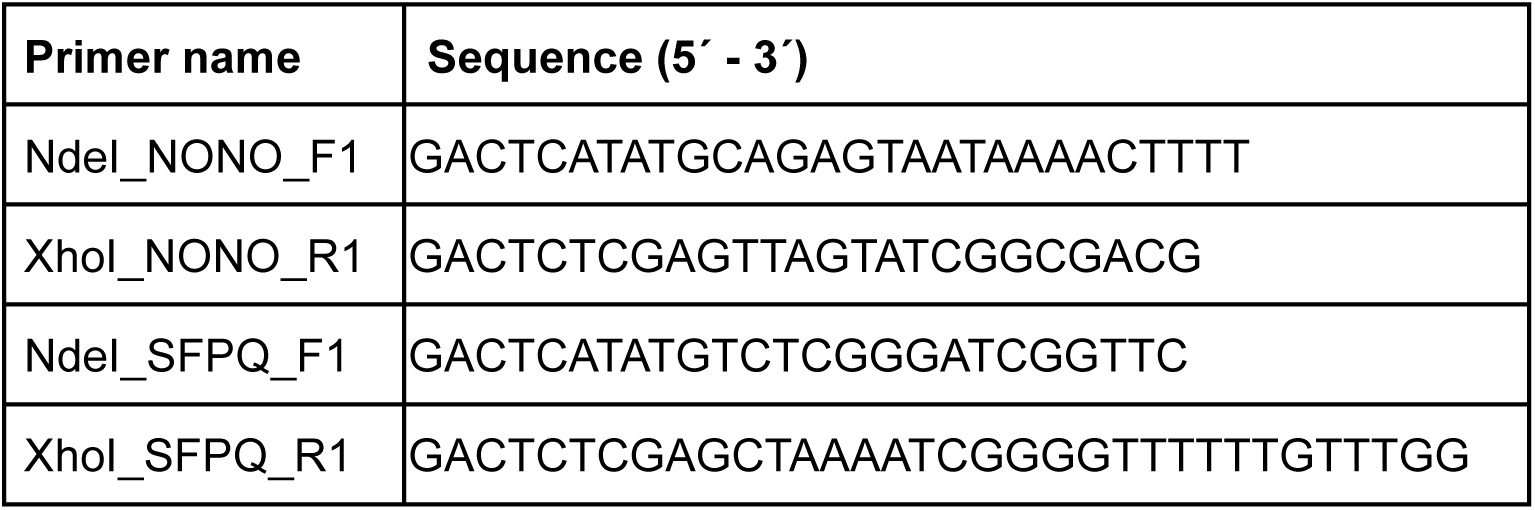

##### Protein purification and labeling

All proteins were expressed as 6x/7xHis-MBP fusion constructs. After transforming bacterial expression plasmids into BL21 *E. coli* (NEB C2527H), cells were grown at 37 °C until reaching OD600 0.6-0.8 and proteins were induced overnight at 18 °C following addition of IPTG (0.5 mM for FUS, NONO, SFPQ, and TDP-43 and 0.4 mM for TEV). Cells were harvested by centrifugation at 14,000 rcf at 4 °C the following day. Cells were resuspended in lysis buffer (described below for each protein) and lysed using probe sonication (QSonica Q500 with 20 kHz converter and 1/4” microtip). Lysate was clarified by centrifugation at 27,000 rcf for 30 min at 4 °C. Following purification, all proteins were verified for purity and size using SDS-PAGE. The following subsections provide purification details for each protein.

#### FUS purification

Lysis buffer consisted of 50 mM HEPES pH 7.4, 1.5 M NaCl, 10% glycerol, 20 mM imidazole, 5 mM β-mercaptoethanol (BME), and EDTA-free protease inhibitor cocktail (Pierce A32965). Clarified lysate was mixed with 0.4-0.5 mL of washed, packed HisPur cobalt resin (Thermo Scientific 89965) per 1 L of cells for 30-60 min at 4 °C. After protein binding, resin was transferred into a gravity flow column and washed with approximately 80-100 resin bed volumes of lysis buffer without protease inhibitor cocktail. Protein was eluted in elution buffer consisting of 50 mM HEPES pH 7.4, 150 mM NaCl, 10% glycerol, 200 mM imidazole, and 5 mM BME. Eluted protein was dialyzed overnight using a 3 mL Slide-A-Lyzer 20K MWCO dialysis cassette (Thermo Scientific 66003) with two rounds of 1 L dialysis buffer consisting of 20 mM HEPES pH 7.4, 150 mM NaCl, 5% glycerol, and 5 mM BME.

#### TDP-43 purification

Lysis buffer consisted of 50 mM HEPES pH 7.4, 1.5 M NaCl, 10% glycerol, 20 mM imidazole, 5 mM BME, and EDTA-free protease inhibitor cocktail. Protein was purified from clarified lysate using an AKTA go fast protein liquid chromatography (FPLC) system (Cytiva). Lysate was loaded onto a 5 mL HisTrap HP column (Cytiva 17524801) equilibrated with lysis buffer, and the column was washed with 10 column volumes of wash buffer consisting of 50 mM HEPES pH 7.4, 1.5 M NaCl, 40 mM imidazole, and 5 mM BME. Protein was eluted with a step to 100% elution buffer consisting of 50 mM HEPES pH 7.4, 1 M NaCl, 500 mM imidazole, and 5 mM BME. Eluted protein was further purified using size exclusion chromatography (SEC) over a Superdex 200 Increase 10/300 GL column (Cytiva 28990944) equilibrated with storage buffer consisting of 20 mM HEPES pH 7.4, 300 mM NaCl, and 5 mM BME.

#### NONO and SFPQ purification

Lysis buffer consisted of 50 mM HEPES pH 7.4, 1.5 M NaCl, 10% glycerol, 20 mM imidazole, 250 mM L-arginine, 0.2% Triton X-100, 5 mM BME, and EDTA-free protease inhibitor cocktail. Proteins were purified from clarified lysates using FPLC following a similar procedure described for TDP-43. Wash buffer consisted of 50 mM HEPES pH 7.4, 1.5 M NaCl, 40 mM imidazole, 250 mM L-arginine, and 5 mM BME. Elution buffer consisted of 50 mM HEPES pH 7.4, 500 mM NaCl, 500 mM imidazole, 250 mM L-arginine, and 5 mM BME. Eluted proteins were further purified using SEC as described for TDP-43. SEC storage buffer consisted of 20 mM HEPES pH 7.4, 500 mM NaCl, and 5 mM BME.

#### TEV protease purification

Lysis buffer consisted of 25 mM HEPES pH 7.4, 300 mM KCl, 10% glycerol, and 20 mM imidazole. HisPur cobalt resin binding and gravity column washing was performed following a similar procedure described for FUS. Resin was further washed using approximately 100 resin bed volumes of lysis buffer containing 10 mM ATP-MgCl_2_, followed by an additional 80-100 resin bed volumes of lysis buffer without ATP-MgCl_2_. Elution buffer consisted of 25 mM HEPES pH 7.4, 150 mM KCl, 10% glycerol, and 250 mM imidazole. Eluted protein was dialyzed overnight using a 3 mL Slide-A-Lyzer 10K MWCO dialysis cassette (Thermo Scientific 66455) with two rounds of 0.5 L dialysis buffer consisting of 25 mM HEPES pH 7.4, 150 mM KCl, 10% glycerol, and 5 mM BME.

#### Protein labeling

Approximately 0.3-0.5 mL of purified proteins were fluorescently labeled with amine-reactive, NHS ester-conjugated Atto 488 or Atto 594 dyes (Sigma 41698-1MG-F and 08741-1MG-F). After resuspending the reactive dye in anhydrous DMSO to a stock concentration of 10 mM, dye was added to the protein at a dye:protein molar ratio of 1:1. The dye was allowed to conjugate to the protein for 20 min at 25 °C, and the mixture was transferred to a 0.5 mL Slide-A-Lyzer 20K MWCO dialysis cassette (Thermo Scientific 66005) and dialyzed with two rounds of 0.5 L dialysis buffer at 4 °C to remove unconjugated dye.

#### Measuring protein and dye concentrations

For FUS, TDP-43, and TEV, protein and conjugated dye concentrations were measured using a Nanodrop spectrophotometer. A260/A280 absorbance ratios were found to be in the range of 0.60-0.65, indicating relatively minor absorbance contributions from residual nucleic acids. For NONO and SFPQ, protein concentrations were measured using the Qubit Protein BR Assay Kit (Invitrogen A50669) according to the manufacturer’s instructions. Small aliquots of labeled and unlabeled proteins were snap frozen in liquid nitrogen and stored at –80 °C.

#### In vitro transcription of NEAT1 RNA fragments

Linear DNA templates for *in vitro* transcription were generated via PCR from respective plasmids using the following forward and reverse primers: 5’-TAATACGACTCACTATAGGGAGTTCTACAGTCCGACGATC-3’ and 5’-GCCTTGGCACCCGAGAATTCCA-3’. *In vitro* transcription was performed using the HiScribe T7 kit (NEB E2040S) following the manufacturer’s instructions using approximately 1 µg of DNA template per 20 µL reaction. 20 U of murine RNase inhibitor (NEB M0314L) was also included in each reaction. To generate fluorescently labeled RNAs, 0.1 µL of Cy5-UTP (Cytiva PA55026) or 1 µL of Atto 488-UTP (Jena Bioscience NU-821-488) were included per 20 µL reaction. Following incubation at 37 °C for 2 h, DNA templates were removed by adding 2 µL of RQ1 RNase-free DNase (Promega M6101) per 20 µL reaction and incubating at 37°C for 20 min. RNA was precipitated by adding LiCl to a final concentration of approximately 830 mM and transferring to –80 °C for at least 15 min. Precipitated RNA was pelleted by centrifugation at 21,300 rcf for 10 min at 4 °C. Supernatant was removed and RNA pellets were washed at least three times with 0.5 mL of RNase-free 80% ethanol. Following resuspension in RNase-free water, RNA concentration was measured using a Nanodrop spectrophotometer and verified for purity and size using a denaturing agarose gel and ssRNA ladder (NEB N0362S). Small aliquots of RNA were stored at –80 °C.

##### Gel electrophoresis mobility shift assays (EMSAs)

20 µL protein-RNA binding reactions were prepared with the following components and allowed to equilibrate at 25 °C for 60 min prior to gel electrophoresis:

● 20 mM HEPES pH 7.4
● 150 mM KCl
● 2 mM MgCl_2_
● 0.05 mM EDTA
● 5% glycerol
● 1 mM TCEP
● 0.1 mg/mL BSA
● 0.1 mg/mL *E. coli* tRNA (Sigma R1753)
● 0.15x purple loading dye (NEB B7024S)
● 1 U/µL RNase inhibitor (RNaseOUT, Invitrogen 10777019 or murine RNase inhibitor, NEB M0314L)
● 2 ng/µL RNA
● variable [protein]

Protein concentrations in EMSAs were as follows:

● FUS and NONO: 25-1600 nM, in two-fold serial dilutions, and 2400 nM
● TDP-43: 25-3200 nM, in two-fold serial dilutions
● SFPQ: 50-800 nM, in two-fold serial dilutions, and 600, 1000 nM

Each EMSA included a reaction without protein. Each SFPQ EMSA contained an additional protein concentration of approximately 1200 nM. However, SFPQ-RNA binding appeared consistently poor at this highest concentration, often with apparently weaker binding than the adjacent lane at lower SFPQ concentration. This effect may have been due to the relatively low concentrations of SFPQ protein stocks, requiring high volumes of un-diluted protein which diluted some components of the EMSA buffer. We therefore omitted RNA bands corresponding to this highest SFPQ concentration from all curves during analysis. SFPQ EMSA gel images displayed in the paper also omit this final, highest SFPQ concentration.

Previous work found that the MBP tag does not substantially impact FUS-RNA affinity^35^, and was therefore not removed to ensure FUS remained soluble. MBP tags were left similarly uncleaved in EMSAs with NONO, SFPQ, and TDP-43.

Prior to preparing binding reactions, a 100 mL, 1% agarose-TBE gel was cast in a 15×10 cm tray with 30-well comb. The set gel was moved to a Wide Mini-Sub Cell GT horizontal electrophoresis system (Bio-Rad), and 15 µL of each binding reaction (30 ng RNA) was loaded per well. The gel was run at 100 V for 90 min at 25 °C in 1x TBE running buffer. The gel was immediately stained in 1x SYBR Gold (Invitrogen S11494), diluted in 1x TBE buffer, with gentle shaking for 2-3 h at 25 °C. A 16.1×12 cm image of the gel was acquired on a Gel Doc imager (Bio-Rad).

Gel band intensities were quantified using ImageJ version 2.14.0. Background subtraction was applied, with a rolling ball radius of 50 pixels, and a rectangular, horizontal region of interest (ROI) was drawn around the unbound RNA bands. The ROI was assigned as the “first lane” in the Gel Analyzer tool, and the Plot Lanes tool was used to generate a plot of band intensities. A vertical line was drawn between each lane in the plot to demarcate bands, and the area under each band was measured using the Wand tool. Band intensities were imported into Matlab 2023b and the protein concentration-dependent fraction of bound RNA, computed

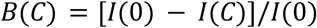

where *I*(*C*) is the protein concentration-dependent band intensity and *I*(0) is the no-protein band intensity. *I*(*C*) values less than 0, corresponding to bands with intensity greater than *I*(0), were filtered out prior to fitting. Individual EMSA experiments, and pooled data from all replicates, were fit to the Hill equation

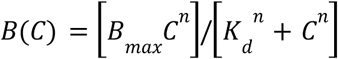

where *B_*max*_* is the maximum fraction of bound RNA at saturation, *K_d_* is the apparent equilibrium dissociation constant, and *n* is the Hill coefficient. Fits were accepted for further analysis if the value of the *K_d_* parameter was greater than the magnitude of its 95% confidence interval from fitting. Accepted *K* estimates were recast in terms of Gibbs free energy upon binding, Δ*G*

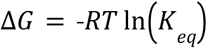

where *R* is the gas constant, *T* is temperature, and *K_*eq*_* is *K_d_* normalized to a reference concentration of 1 M. ΔΔ*G* was then computed by subtracting Δ*G* from a reference value, chosen as the overall median Δ*G* of FUS binding to C1, C2, and C3 RNAs.

#### Fluorescence microscopy

Imaging and FRAP was performed on a Zeiss LSM 980 laser scanning confocal microscope equipped with Airyscan 2 detector, 32-channel GaAsP array detector, 405/488/561/639 nm lasers, 63x/1.4 NA oil immersion and 40x/1.2 NA water immersion objectives, and HP Z6 workstation with Zeiss ZEN 3.7 software.

#### In vitro condensate experiments

Prior to preparing mixtures for condensate experiments, wells of 96-well glass bottom plates (Greiner Bio-One 655892) were passivated with 1 wt/vol% Pluronic F-127 for at least 30 min at 25 °C^57^. Wells were subsequently washed at least 10 times with ultrapure water and dried in a 50 °C oven for at least 20 min.

Proteins and RNAs were diluted in droplet buffer consisting of 20 mM HEPES pH 7.4, 150 mM NaCl, and 2 mM TCEP. Final protein and RNA concentrations used in experiments are indicated in the figure legends. The final volume of each mixture was 100 µL. Prior to diluting NONO in droplet buffer, the protein stock, stored in buffer containing 500 mM NaCl, was pre-diluted in low-salt buffer (20 mM HEPES pH 7.4, 10 mM NaCl, and 2 mM TCEP) such that the final NaCl concentration was 150 mM. For experiments with FUS and TDP-43, Atto 488 or Atto 594-labeled protein was included such that 5% of protein was dye-labeled; for NONO, Atto 594-labeled protein was included such that 0.5-1% of protein was dye-labeled. TEV protease was added last to each mixture, to a final concentration of 1.5 µM, to cleave the MBP tag and initiate condensation. Immediately after TEV addition, solutions were gently mixed, loaded into passivated wells, and incubated for 3 h at 25 °C. Z-stacks of condensates were acquired with a z-spacing of 0.24 µm. At least three z-stacks were acquired per experimental replicate. Images were acquired using identical imaging settings (e.g. laser power, pixel dwell time, detector gain) across different RNAs and experimental replicates in order to facilitate pooling of replicate data and enable direct comparison of dense phase intensity measurements among different experimental conditions.

#### Image analysis

Images were analyzed using custom Python scripts. Z-stacks were imported and the frame with the greatest mean brightness was chosen for analysis. An automatic threshold was applied using the Otsu method, and the resulting binary mask was used to compute droplet properties using methods.regionprops_table from scikit-image version 0.25.0. When analyzing two-color images, the following channels were chosen as reference to generate the mask, which was subsequently applied to both channels for analysis:

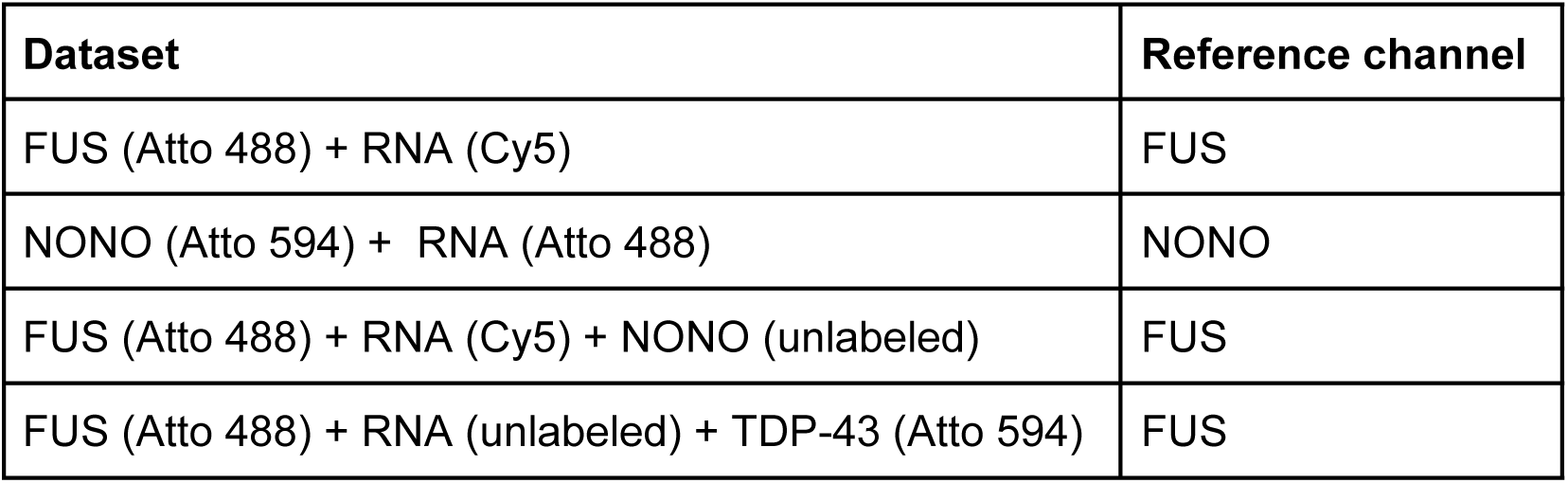

| Dataset | Reference channel |
| --- | --- |
| FUS (Atto 488) + RNA (Cy5) | FUS |
| NONO (Atto 594) + RNA (Atto 488) | NONO |
| FUS (Atto 488) + RNA (Cy5) + NONO (unlabeled) | FUS |
| FUS (Atto 488) + RNA (unlabeled) + TDP-43 (Atto 594) | FUS |

During analysis, the mean intensities of smaller droplets were found to scale with droplet area, indicating that lower intensity pixels near droplet edges (corresponding to the transition zone between dense and dilute phases) disproportionately skewed mean intensity estimates toward lower values for smaller droplets compared to larger droplets. To minimize this size-dependent intensity effect, the presented data represent centroid intensity estimates for the 50 largest droplets per experimental replicate. Centroid intensities were computed as the average of five pixels at the estimated centroid position of the droplet: the pixel nearest to the centroid and the four pixels directly bordering it.

Because each 1000 nt NEAT1 fragment contained different fractions of uridine (U), dye:RNA labeling ratios were unequal across RNAs owing to variable incorporation of labeled U during *in vitro* transcription. To facilitate relative comparison of dense phase RNA intensities across different RNAs, a linear scaling factor for each RNA was required to adjust for labeling differences. Dye concentrations for many of our RNA stocks were too dilute to confidently estimate labeling ratios from spectrophotometry, hindering our ability to compute scaling factors directly from labeling ratios. However, we found that with several other, more highly concentrated RNA stocks, the labeling ratio consistently scaled linearly with U fraction (Fig. S3E). Thus, U fraction can serve as a proxy for labeling ratio when computing scaling factors. Specifically, scaling factors were computed as the ratio of the E1 U fraction relative to that of the RNA of interest, thereby rescaling all RNA intensities relative to E1. In order to facilitate visual comparison of RNA dense phase intensities, these same scaling factors were also used to appropriately rescale the upper contrast bounds of RNA images displayed in Fig. 2D.

#### Fluorescence recovery after photobleaching (FRAP)

Single points within droplets were bleached using 488 and 561 nm lasers and images were acquired at fixed z positions every three s, with three frames acquired prior to bleaching. Bleached droplets were automatically detected and time-dependent average intensities at droplet centroids were computed using a custom Python script. Briefly, the first frame of each time series was automatically thresholded using the Otsu method, and the resulting binary mask was applied to each frame to compute time-dependent droplet properties using methods.regionprops_table from scikit-image version 0.25.0. Centroid intensities were computed as described in the “Image analysis” section. Bleached droplets were defined as having post-bleach centroid intensities less than 25% that of pre-bleach in the prior frame.

After computing the time-dependent intensity at droplet centroids, *C*(*t*), the normalized, time-dependent intensity, *C_*norm*_* (*t*), was calculated as described in ref^58^

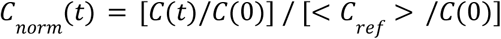

where *C*(0) is the centroid intensity immediately post-bleaching and <**C*_*ref*_*> is the average, time-dependent intensity at the centroids of unbleached droplets. Unbleached droplets were selected for use in computing <**C*_*ref*_*> if the initial centroid intensity was within 20% that of the bleached droplets. *C*_*norm*_ (*t*) for individual droplets and for average data from all droplets was fit to the 2D, single-component FRAP recovery model^44^

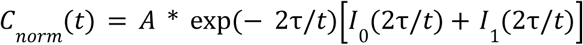

where *A* is the mobile fraction, τ is the recovery time constant, and *I*_0_ and *I*_1_ are zero and first order modified Bessel functions. Fitting was performed in Python using the curve_fit module from scipy.optimize version 1.14.1.

#### Biological materials

All unique biological materials generated for this study are available from the corresponding author upon request.

#### Statistics and reproducibility

Sample sizes were not predetermined. All microscopy images are representative of experimental replicates that were repeated independently on separate days at least three times. No data were excluded during analysis. Blinding and randomization was not performed, as all microscopy data were acquired using identical imaging settings and analysed using automated analysis routines.

## Supplementary Figures

**Fig. S1.**
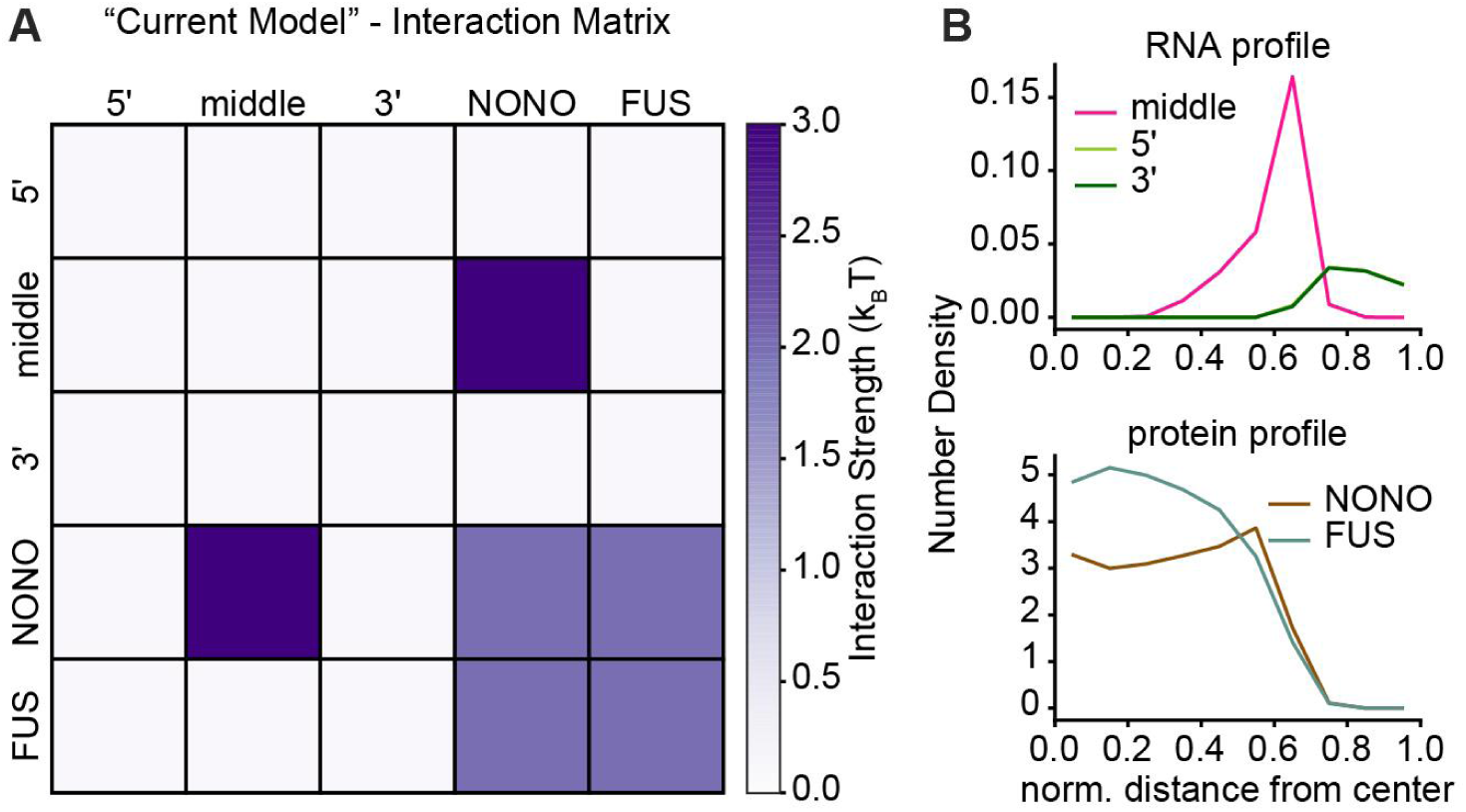
Simulating the current model of paraspeckle assembly. (**A**) Matrix of interactions used in simulations. (**B**) Radial density profiles of NEAT1 segments (top) and proteins (bottom). Plots represent average profiles of the largest clusters from five simulations.

**Fig. S2.**
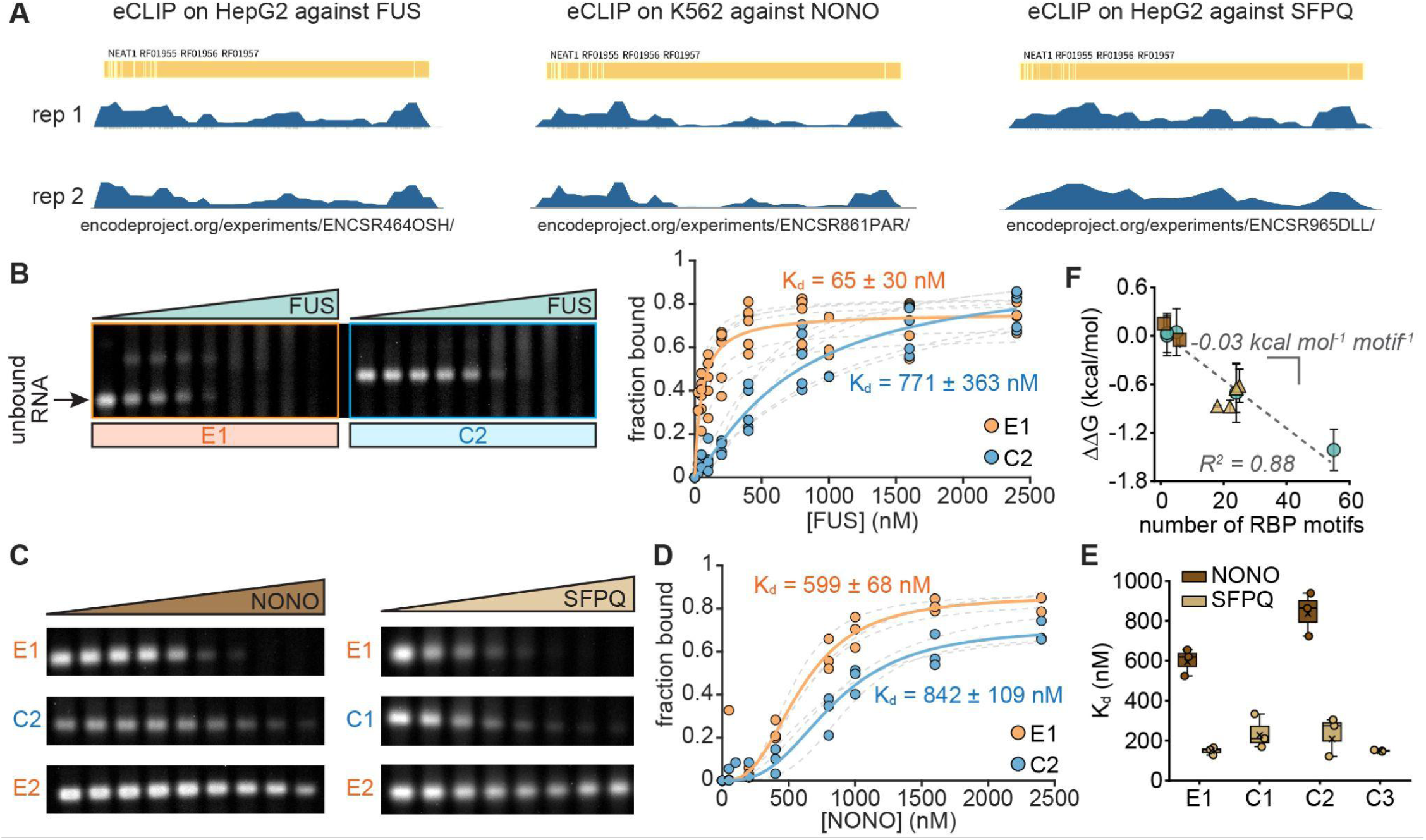
Paraspeckle core RBPs display unexpected binding to NEAT1. (**A**) NEAT1 eCLIP data from ENCODE for the indicated cell type against the indicated protein. (**B**) Background-subtracted image of unbound RNA in FUS EMSA gel with E1 and C2 fragments (separate replicate from the one displayed in Fig. 2B). Plot: Fraction of bound E1 or C2 RNA as a function of FUS concentration. Gray dashed curves and solid curves in color indicate Langmuir isotherm fits to individual replicates and pooled data from all replicates, respectively. (**C**) Background-subtracted images of unbound RNA in EMSA gels with the indicated proteins and RNAs. For each protein, the three sets of lanes were cropped from the same gel, with equal contrast settings applied. Poor NONO and SFPQ binding to E2 hindered K_d_ estimation. (**D**) Fraction of bound E1 or C2 RNA as a function of NONO concentration, formatted similar to (B). (**E**) NONO-RNA and SFPQ-RNA K_d_ estimates from EMSAs. Circles and x’s indicate K_d_ estimates from individual replicates and pooled data, respectively. Boxes indicate IQRs with medians as bisecting lines and whiskers as 1.5*IQR. (**F**) Data from Fig. 2C, expressed as ΔΔG, as a function of motif abundance. ΔΔG values were computed relative to the overall median of FUS binding to C1, C2, and C3. Dashed gray line indicates linear regression.

**Fig. S3.**
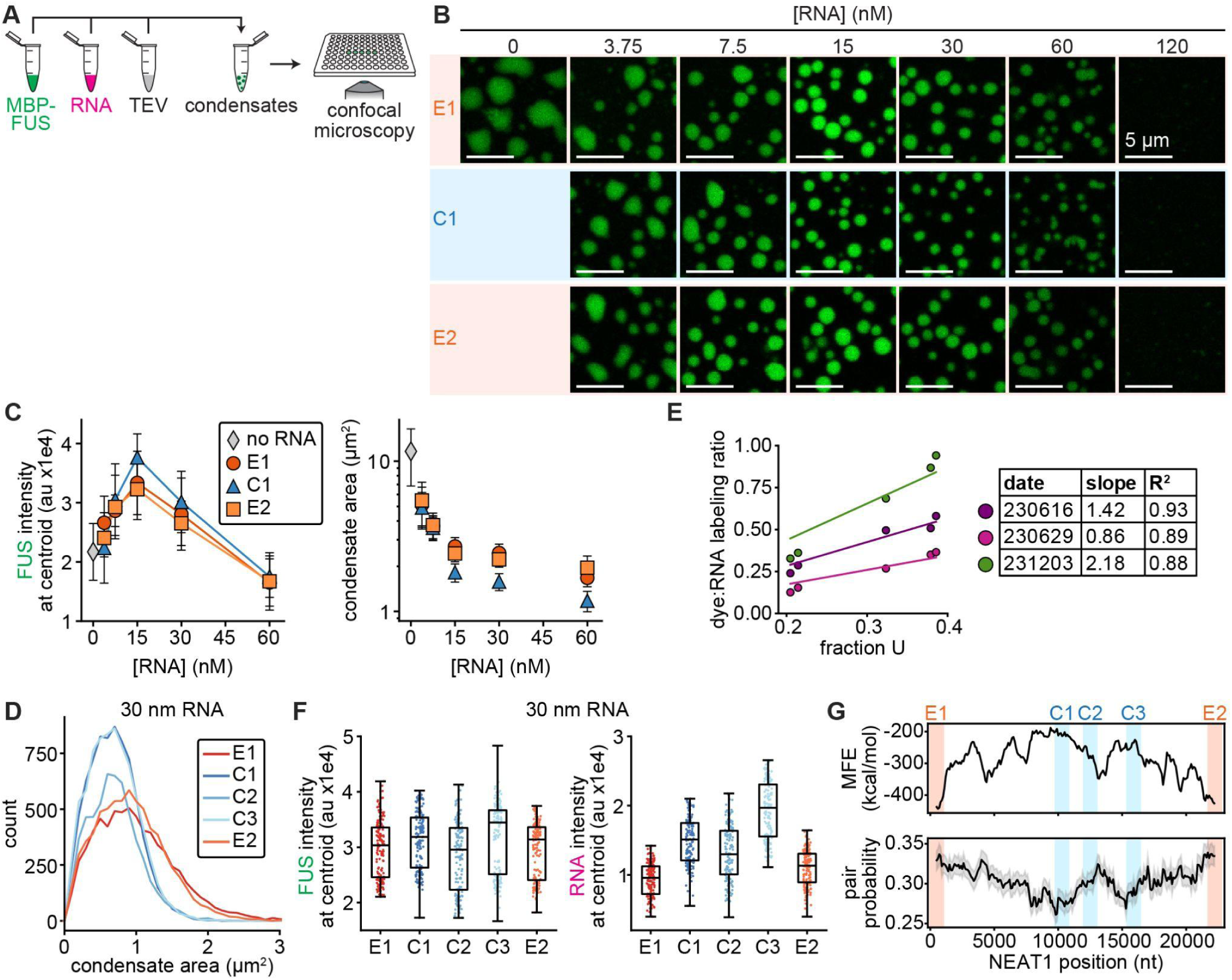
FUS condensates assembled with middle region RNAs are smaller owing to stronger RNA-RNA interactions in the dense phase. (**A**) Schematic of *in vitro* condensation experiments. (**B**) Confocal slices of condensates assembled with 2 µM FUS (green) and RNA at the indicated concentrations after 3 h at 25 °C. FUS channel is contrasted equally in all images. (**C**) FUS intensity at condensate centroids (left) and condensate area (right) as a function of RNA concentration for the three indicated RNAs. Data points and error bars indicate mean ± one standard deviation for the 50 largest condensates in each experimental replicate. Note that at RNA concentrations of 15 nM and higher, condensate area is consistently lower for C1 compared to E1 and E2. (**D**) Histograms of condensate area, assembled with 2 µM FUS and 30 nM RNA. (**E**) Dye:RNA labeling ratio as a function of uridine (U) fraction for the five RNAs E1, C1, C2, C3, and E2. Data represent three RNA preps from different days, each using the same reaction master mix for the five RNAs. Lines indicate linear regression with y-intercept set to 0. Table summarizes fit slopes and R^2^ values. (**F**) Boxplots of FUS (left) and RNA (right) intensity at condensate centroids for condensates assembled with 2 µM FUS and 30 nM RNA. Data represent the 50 largest condensates from each experimental replicate. Circles indicate individual condensates, boxes indicate IQR with medians as bisecting lines and whiskers as 1.5*IQR. (**G**) Minimum free energy (MFE) of the optimal secondary structure (top) and base pair probability (bottom) within 1000 nt windows of NEAT1, tiled every 100 nt from 5’ to 3’. Bottom plot indicates the average pair probability of all nt’s within each window, shaded error bar indicates standard error of the mean. The higher MFE values and lower pair probabilities of middle region RNAs suggest that the middle region is less structured and more single-stranded. RNA structure prediction was performed using the ViennaRNA package for Python (Methods). Orange and blue vertical bars indicate the five NEAT1 fragments used in this study.

**Fig. S4.**
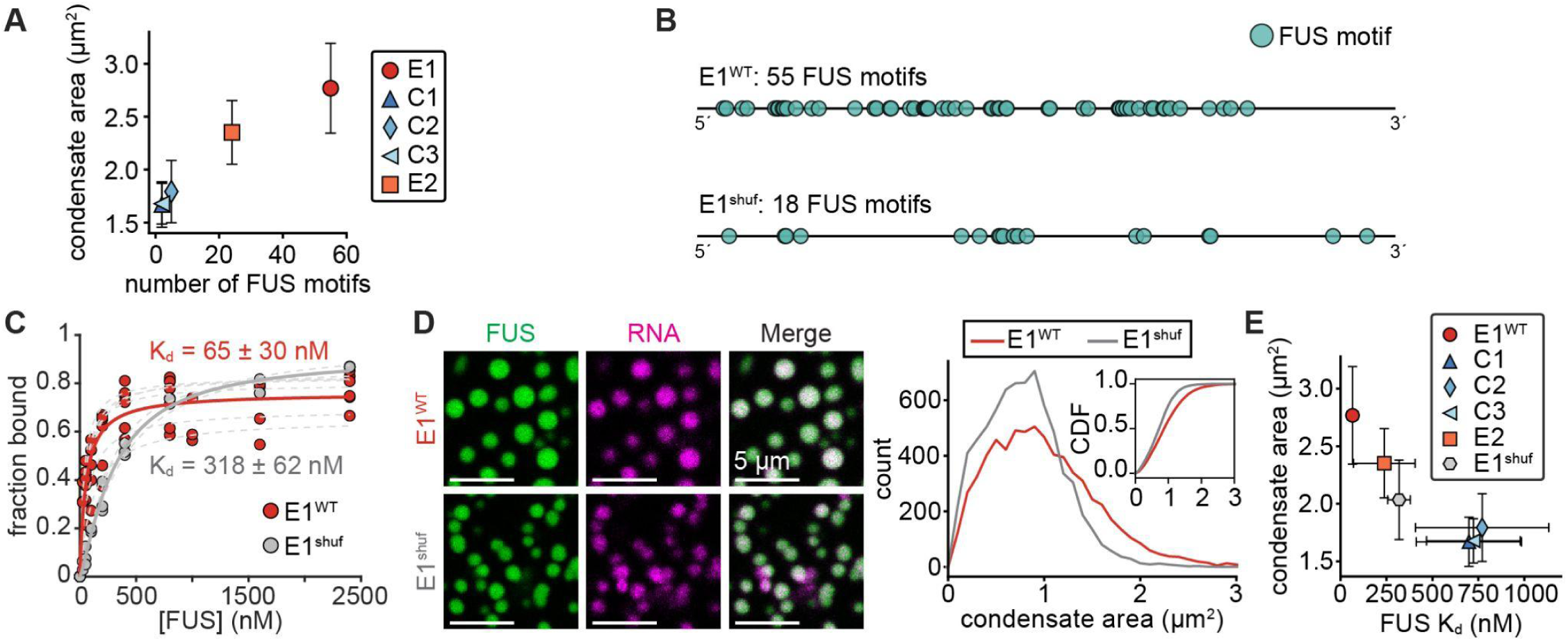
Condensate area scales with FUS-NEAT1 affinity. (**A**) Condensate area as a function of the number of FUS motifs within each RNA. (**B**) Schematic of FUS motifs in E1^WT^ and a shuffled E1 sequence, E1^shuf^. (**C**) Fraction of bound E1^WT^ or E1^shuf^ RNA as a function of FUS concentration from EMSAs. Gray dashed curves indicate Langmuir isotherm fits to individual replicates, solid curves in color indicate fits to pooled data from all replicates. E1^WT^ data repeated from Fig. S2B. (**D**) Left: Confocal slices of condensates assembled with 2 µM FUS (green) and 30 nM E1^WT^ or E1^shuf^ RNA (magenta). FUS and RNA contrasted equally. Right: Histograms of condensate area from experiments corresponding to the images on the left. (**E**) Condensate area as a function of FUS-RNA binding affinity, K_d_. Data points and error bars in (A) and (E) represent mean ± one standard deviation for the 50 largest condensates in each experimental replicate.

**Fig. S5.**
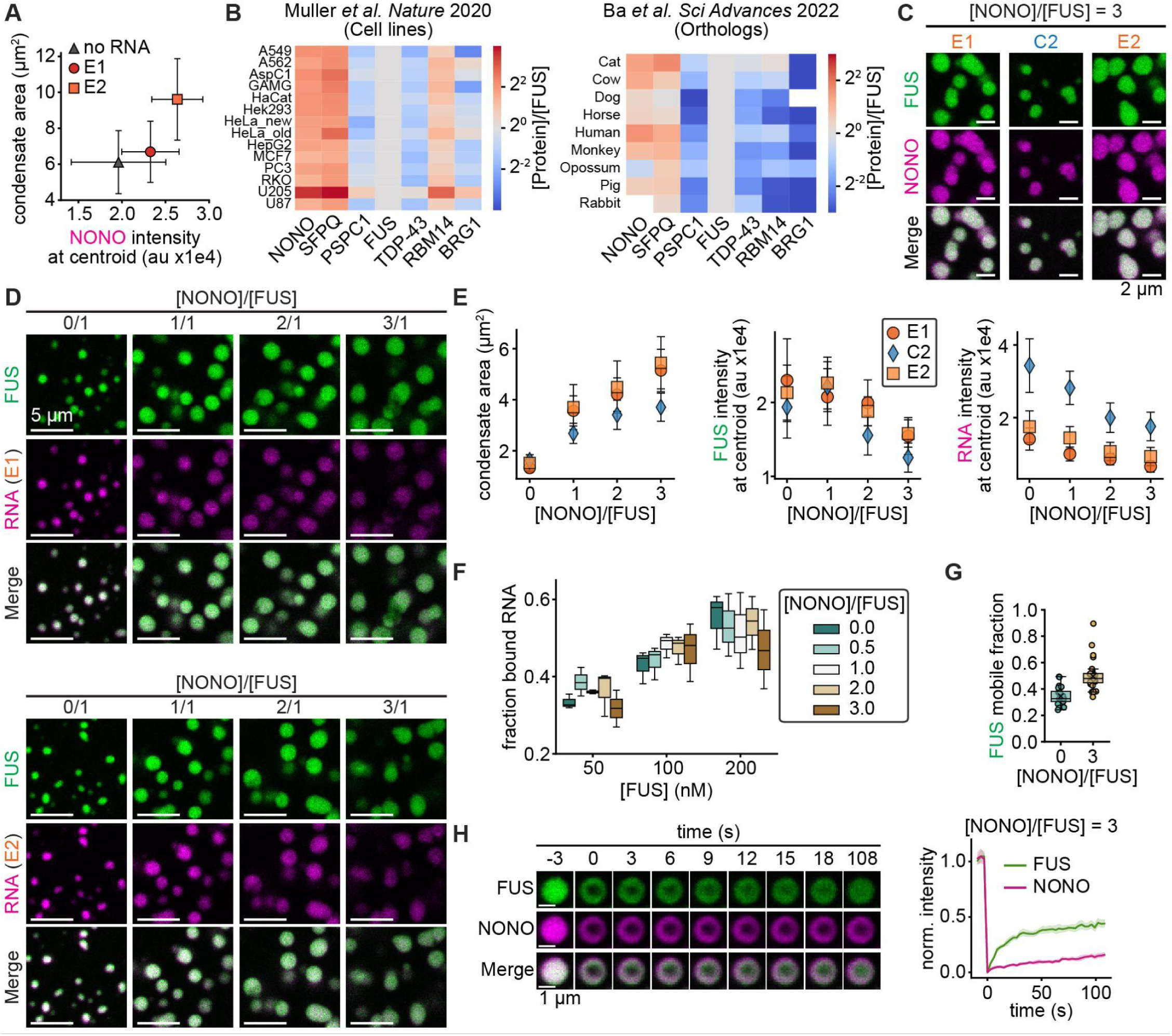
NONO/FUS stoichiometry rewires the properties of FUS-RNA condensates. (**A**) Condensate area as a function of NONO intensity at condensate centroids for condensates assembled with 2 µM NONO and 15 nM of the indicated RNA. (**B**) Heat maps of paraspeckle protein abundance relative to FUS in the indicated cell lines (left, data from ref^42^) and in fibroblasts from the indicated mammalian species (right, data from ref^43^). (**C**) Confocal slices of condensates assembled with 1 µM FUS (green) + 15 nM of the indicated RNA (unlabeled) + 3 µM NONO (magenta) after 3 h at 25 °C. FUS and NONO channels each contrasted equally in all images. (**D**) Confocal slices of condensates assembled with 1 µM FUS (green) + 15 nM of the indicated RNA (magenta) + NONO (unlabeled) at the indicated concentrations after 3 h at 25 °C. FUS and RNA channels each contrasted equally in all images. (**E**) Condensate area (left) and FUS (middle) and RNA (right) intensity at condensate centroids as a function of NONO/FUS stoichiometry with the indicated RNAs, from experiments corresponding to (D). Data points and error bars in (A, E) indicate mean ± one standard deviation for the 50 largest condensates from each experimental replicate. (**F**) Fraction of bound E1 RNA from EMSAs performed with the indicated FUS concentrations in the presence of the indicated NONO/FUS stoichiometries. (**G**) FUS mobile fraction estimates from FRAP fits. Circles and x’s indicate estimates from fits to individual droplets and average data, respectively. Boxes in (F, G) indicate IQR with medians as bisecting lines and whiskers as 1.5*IQR. (**H**) Confocal slices of FUS (green) and NONO (magenta) photobleaching recovery within droplets assembled with 1 µM FUS + 15 nM E2 RNA + 3 µM NONO. Plot shows normalized FUS and NONO intensity recovery profiles as a function of time for experiments corresponding to the images on the left. FUS profile repeated from Fig. 3F. Curves and shaded error bars indicate average ± 95% CI.

**Fig. S6.**
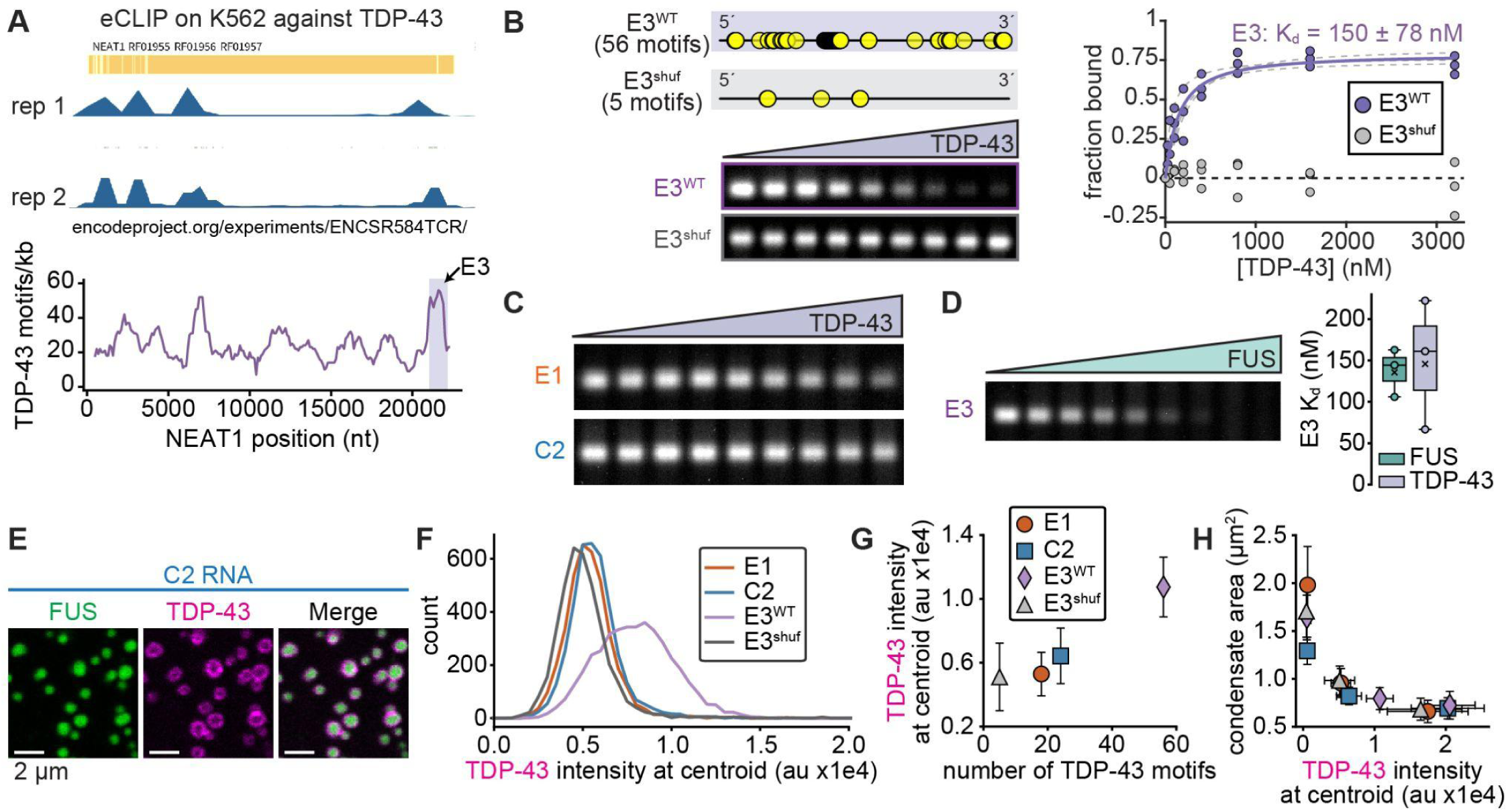
TDP-43 binds abundant cognate motifs in a 3’-proximal NEAT1 fragment to oppose FUS condensation. (**A**) Top: NEAT1 eCLIP data from ENCODE for K562 cells against TDP-43. Bottom: the number of TDP-43 motifs within 1000 nt windows of NEAT1, tiled every 100 nt from 5’ to 3’. Plot repeated from Fig. 4A. Purple vertical bar indicates the E3 fragment. (**B**) Schematic of TDP-43 motifs in E3^WT^ and a shuffled E3 sequence, E3^shuf^. Below: background-subtracted image of un-bound E3^WT^ and E3^shuf^ RNA in a TDP-43 EMSA gel. Lanes cropped from the same gel, with equal contrast settings applied. Plot: Fraction of bound E3^WT^ or E3^shuf^ RNA as a function of TDP-43 concentration. Gray dashed curves and solid curve in color indicate Langmuir isotherm fits to individual replicates and pooled data from all replicates, respectively. Undetectable TDP-43 binding to E3^shuf^ over the tested concentration range hindered Langmuir isotherm fitting. (**C**) Background-subtracted images of unbound RNA in TDP-43 EMSA gels for E1 and C2 RNAs. Each set of lanes was cropped from a different gel, but equal contrast settings were applied to both images. Poor TDP-43 binding to E1 and C2 hindered K_d_ estimation. (**D**) Left: background-subtracted image of unbound RNA in FUS EMSA gel for E3^WT^. Right: FUS and TDP-43 K_d_ estimates for E3^WT^. Circles and x’s indicate K_d_ estimates from individual replicates and pooled data, respectively. Boxes indicate IQRs with medians as bisecting lines and whiskers as 1.5*IQR. (**E**) Confocal slices of condensates co-assembled with 2 µM FUS (green) + 0.2 µM TDP-43 (magenta) + 15 nM of unlabeled C2 RNA after 3 h at 25 °C. (**F**) Histograms of TDP-43 intensity at the centroids of condensates from experiments corresponding to (E) and Fig. 4B. (**G**) TDP-43 intensity at condensate centroids, from experiments corresponding to (E) and Fig. 4B, plotted as a function of the number of TDP-43 motifs per RNA. (**H**) Condensate area as a function of TDP-43 intensity at condensate centroids. Data are from experiments in which TDP-43 was added at concentrations of 0, 0.2, and 0.5 µM, with fixed FUS and RNA concentrations indicated in (E). Data points and error bars in (G, H) indicate mean ± one standard deviation for the 50 largest condensates from each experimental replicate (Methods).

**Fig. S7.**
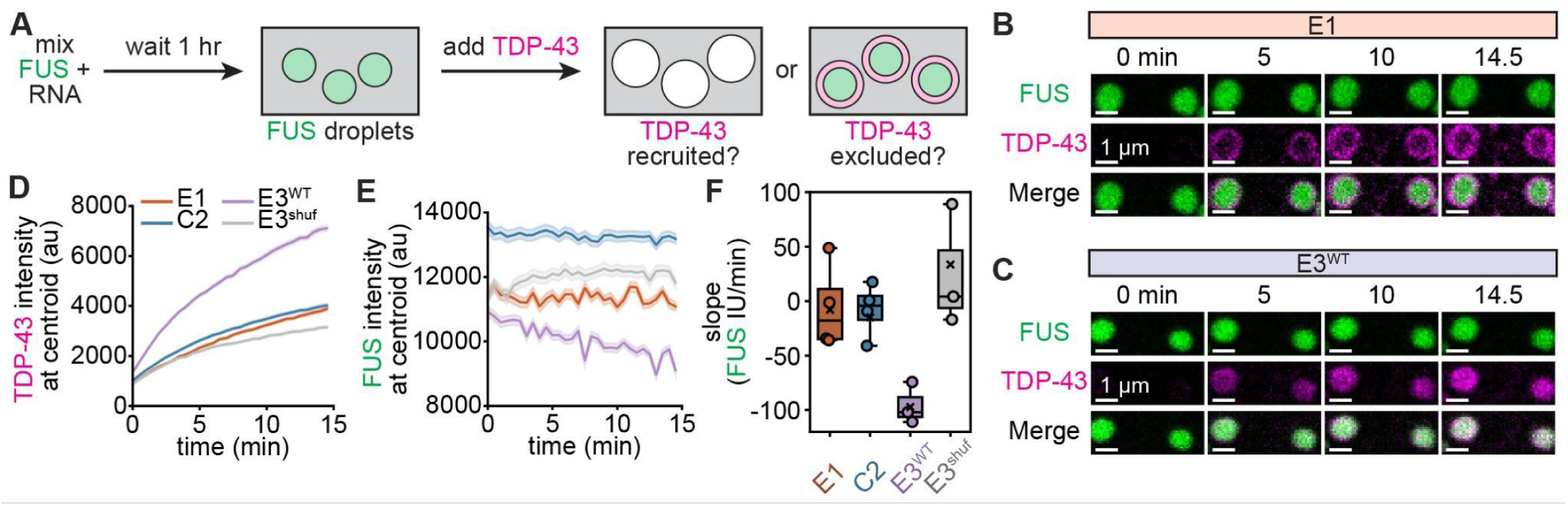
TDP-43 evicts FUS from pre-assembled condensates with E3^WT^ RNA. (**A**) Workflow for examining TDP-43 recruitment to pre-assembled FUS + NEAT1 condensates. (**B, C**) Confocal time series of condensates pre-assembled with 2 µM FUS (green) + 15 nM of either E1 (B) or E3^WT^ (C) RNAs following addition of 2 µM TDP-43 (magenta) at the indicated time points. (**D, E**) TDP-43 (D) and FUS (E) intensity at the centroids of FUS condensates assembled with the indicated RNAs, plotted as a function of time after TDP-43 addition. Curves and shaded error bars indicate average ± 95% CI. (**F**) Slope of FUS intensity over time following addition of TDP-43 (IU: intensity units), quantified from data in (E). Circles and x’s indicate slopes from fits to individual replicates and pooled data, respectively. Boxes indicate IQR with medians as bisecting lines and whiskers as 1.5*IQR. The more negative slope for E3^WT^ indicates that FUS was evicted from condensates by TDP-43 more strongly compared to other RNAs.

**Fig. S8.**
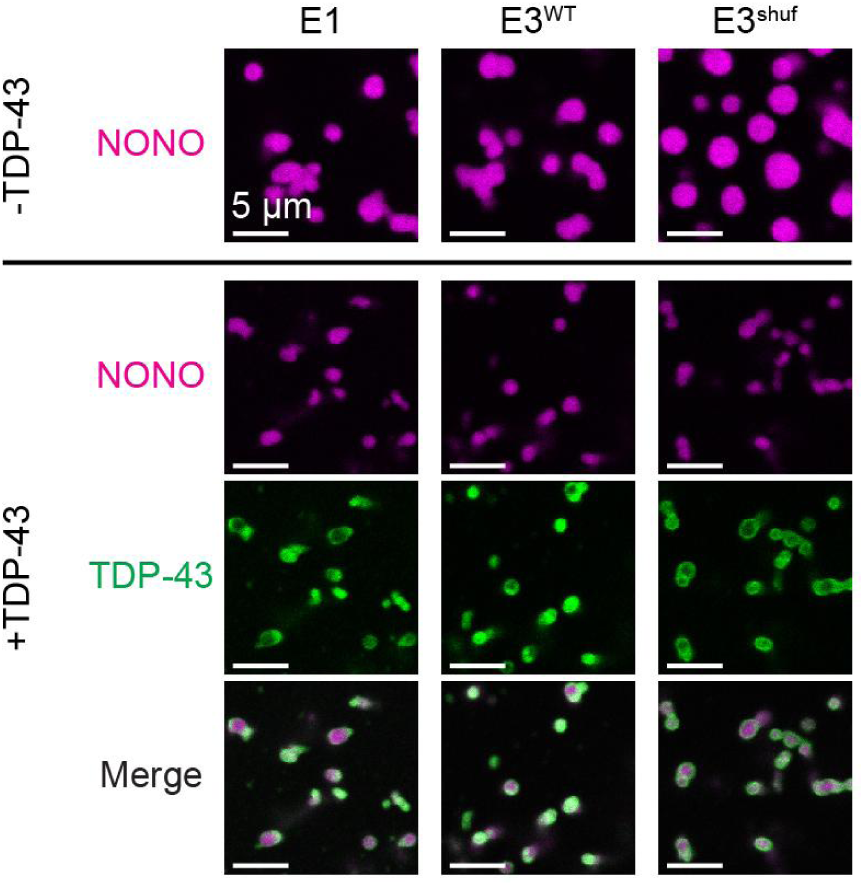
TDP-43 opposes NONO condensation by forming surfactant-like layers on condensate surfaces. Confocal slices of condensates assembled with 2 µM NONO (magenta) + 15 nM of the indicated RNAs ± 0.2 µM TDP-43 (green) after 3 h at 25 °C. NONO and TDP-43 channels are each contrasted equally across all images.

**Fig. S9.**
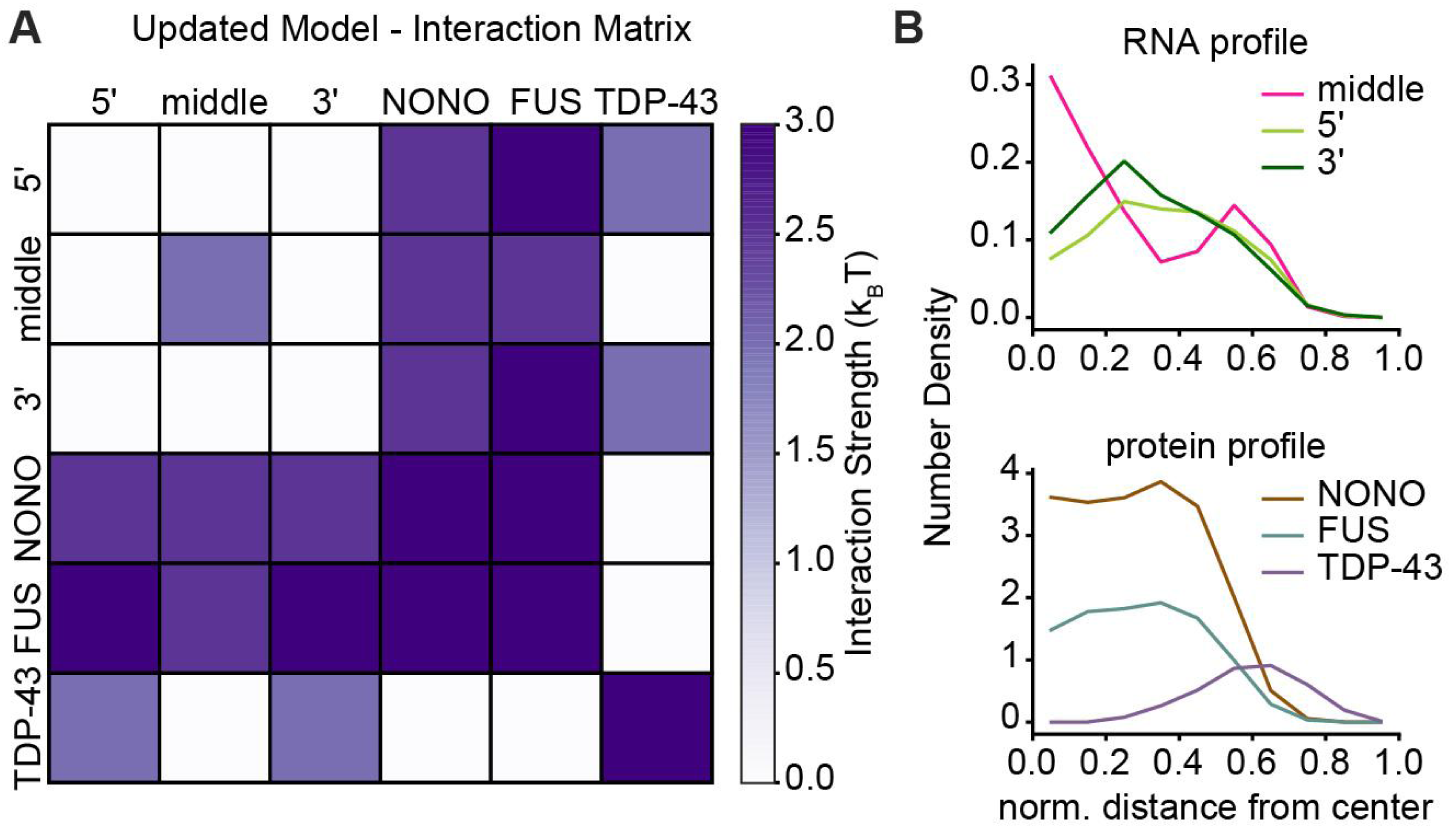
An updated model relates the observed network of biomolecular interactions to paraspeckle assembly. (**A**) Matrix of interactions used in simulations. (**B**) Radial density profiles of NEAT1 segments (top) and proteins (bottom). Plots represent average profiles of the largest clusters from five simulations.

**Fig. S10.**
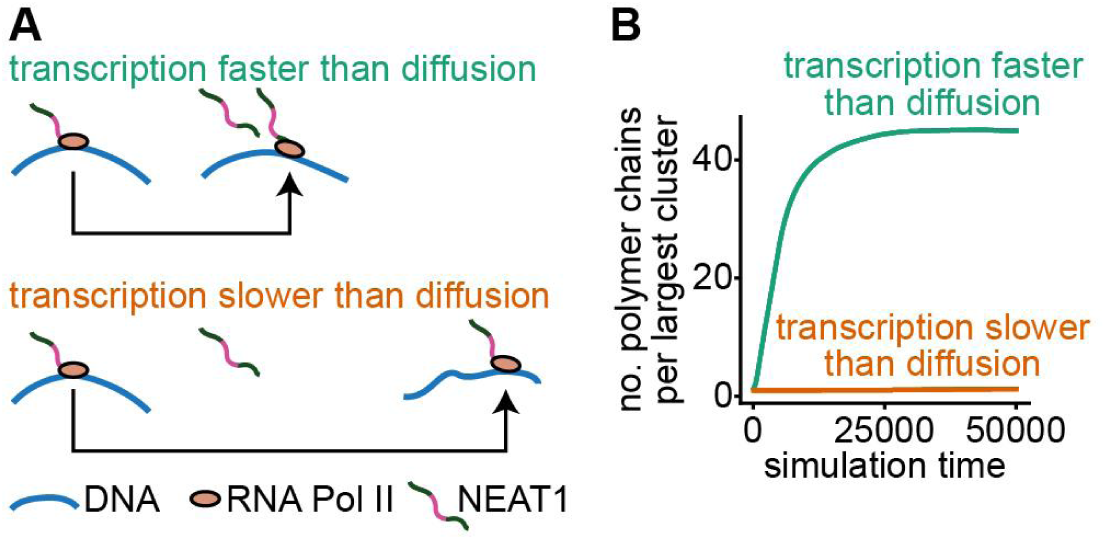
Transcriptional dynamics determine the number of NEAT1 transcripts incorporated into clusters. (**A**) Schematic of the two model scenarios. Upper: when transcription is faster than chromatin diffusion (i.e. burst transcription), multiple transcripts accumulate within close spatial proximity to each other. Lower: when transcription is slower than chromatin diffusion (i.e. latent or paused transcription), transcripts disperse over a relatively large volume. (**B**) The number of NEAT1 polymer chains within the largest cluster in simulations as a function of time. See Methods for simulation details.

## Supplementary Table Legend

**Table S1.** Interaction parameters and key model input values used in coarse-grained simulations of the current model (Fig. 1 and S1A) and updated model (Fig. 5 and S9A) of paraspeckle assembly.

